# Uncovering the molecular basis of kinase activity and substrate recognition with phospho-PCA

**DOI:** 10.1101/2025.11.07.687244

**Authors:** Changhua Yu, Edward Pao, David Van Valen

**Affiliations:** Division of Biology and Biological Engineering, California Institute of Technology, Pasadena, CA 91125, USA; Howard Hughes Medical Institute, Chevy Chase, MD 20815, USA

**Keywords:** protein kinases, deep mutational scanning, substrate specificity, phosphorylation, kinase activity, allostery, phospho-PCA, protein complementation assay

## Abstract

Protein kinases relay information to various cellular processes, and their dysregulation underlies numerous human diseases. Despite their importance, our understanding of how kinase domains and their variants impact protein stability, catalytic activity, and substrate recognition is incomplete. In this work, we develop the phosphorylation protein complementation assay (phospho-PCA), which enables quantitative measurements of kinase-substrate interactions by coupling them to the growth of budding yeast, thereby enabling deep mutational scanning of kinase domains. When combined with deep mutational scans targeting folding stability and Bayesian modeling, phospho-PCA can disentangle the relative impact of mutations on kinase domain stability, catalytic activity, and substrate specificity. We demonstrate the accuracy and breadth of phospho-PCA, showing its applicability to both tyrosine and serine/threonine kinase domains. We then apply our method to three closely related protein kinases with distinct substrate preferences, evaluating over 15,000 kinase variants against a panel of three substrates for both catalytic activity and substrate specificity. The resulting dataset constitutes the largest and most detailed variant-to-function map assembled for this enzyme family to date, revealing numerous mutations that alter kinase activity and substrate specificity. Physics-based modeling reveals how these mutations operate through diverse mechanisms, including long-range allosteric communication, to alter both activity and substrate specificity. Given its scalability, we believe phospho-PCA can measure the functional impact of variants across the entire human kinome.

## INTRODUCTION

In response to upstream signals, protein kinases selectively transfer phosphate groups to their substrates, significantly affecting the behavior of the substrate protein by causing changes in enzymatic activity, cellular localization, and affinity to binding partners^1^.^2,3^. Given the central role that protein kinases play in the cell’s information processing capabilities, their ability to control their activity and select a specific set of substrates is essential to their function. Dysregulation of protein kinases can be caused by single missense mutations, or variants, and these variants are important drivers of numerous diseases^4–7^, including autoimmunity and cancer. Improved knowledge of how variants affect catalytic activity of kinase domains as well as substrate recognition will be critical for understanding broader concepts of protein stability, structural determinants of specificity, and mechanisms of allostery, with the potential to generate potent and specific molecules that alter kinase activity without off-target effects^8–10^.

A key gap in our knowledge is a mechanistic understanding of how kinase domains contribute to substrate specificity. While physical co-localization of kinases with their substrate, often mediated by interactions between peptide binding domains and short linear interaction motifs, is a key component of substrate specificity^11^, a growing body of work demonstrates that the kinase domain itself makes a significant contribution to substrate recognition^12^. First, domain swapping experiments on receptor tyrosine kinases have shown that cells expressing chimeric receptors will develop a phosphorylation profile consistent with the substrate preference of the kinase domain rather than the base receptor^13–18^. Complementing this work are studies on RET^19,20^, CAMK2^21^, c-Src^22,23^, Kit^24^, PKA^25,26^, HER2^27^, and ERK^28^, which show that single-point mutations in kinase domains can change their substrate preferences and, in some cases, drive pathogenicity^20,24–27^. Second, while kinase domains exhibit significant structural similarity, they display substantial variations in the charge and hydrophobicity of their surface residues^29,30^. This observation, in combination with evolutionary analyses of tyrosine kinases, supports the hypothesis that kinase domains originated from a promiscuous common ancestor and evolved to develop distinct and diverse substrate preferences^31^.

Despite significant advances in profiling kinase-substrate interactions, the field has been limited by available methods^22,32–36^. For example, peptide positional scanning libraries have generated consensus motifs for protein kinases at scale^35–37^, as well as insights about substrate specificity^22^. However, these methods are unable to assess kinase variants in parallel, making it challenging to use them to study the impact of variants on substrate specificity. Despite an explosion of data on protein structures in recent years, few structures exist of kinase domains in complex with their substrates^38^. Deep mutational scanning is another technology well-suited for understanding the impact of variants on protein function^39,40^. With deep mutational scanning, the impact of thousands of mutations is measured in parallel by coupling protein activity to cell growth. Provided that each variant is suitably barcoded, this coupling enables quantification by using next-generation sequencing to measure the abundance of each variant and, therefore, its activity. Deep mutational scanning has been applied to various systems, including viral fitness^41–43^, photosynthetic enzymes^44^, G-proteincoupled receptors^45,46^, protein-protein interactions^47,48^, as well as kinases such as ABL^49^, MET^50^, EGFR^51^, Syk^52^, ERK2^53^, TYK2^54^, and c-Src^55,56^. One limitation of these data for protein kinases is that most do not distinguish between increased protein abundance and kinase activity, with the exception of two recent studies^54,56^. Critically, these prior studies measure overall kinase activity and are unable to reveal changes in substrate preferences among kinase variants.

In this work, we were able to generate the first comprehensive maps of how mutations within the kinase domain impact both catalytic activity and substrate recognition. We focus on Fyn, Lck, and c-Src of the Src family kinases (SFKs). These three non-receptor tyrosine kinases share ≈70–80% sequence similarity but play distinct roles in regulating differentiation, immune responses, and cell growth. Fyn has a broad tissue distribution and is involved in T-cell signaling^57–59^ and neuronal development^60,61^; it is currently being investigated as a target for neurodegenerative diseases^62,63^. Lck is expressed in T-cells and is a key component of the T-cell activation signaling cascade^64–67^. c-Src is one of the most well-studied oncogenes; it is widely expressed and regulates cell adhesion, proliferation, and survival^68^. The different roles these kinases have assumed are partly due to their varying substrate preferences. Variants of SFKs have been linked to numerous clinical conditions, including immunodeficiency and cancer^69–74^.

Here, we develop the phosphorylation protein complementation assay (phospho-PCA), which uses a growth-based biosensor to quantitatively couple kinase-substrate interactions to the growth of budding yeast. We demonstrate that phospho-PCA enables deep mutational scanning of kinase domains, using the interaction strength with a specific substrate as the readout. Furthermore, we demonstrate that combining mutational scans from phospho-PCA and abundance-PCA, which measures folding stability of protein variants, with statistical modeling enables us to quantify the relative impact of mutations on kinase domain stability, kinase activity, and substrate preference. We perform mutational scans with abundance-PCA and phospho-PCA of Fyn, Lck, and c-Src kinase domains, measuring the impact of ≈ 5000 single amino acid variants (≈ 98% coverage) for each kinase when paired with representative Fyn, Lck, and c-Src substrates. We demonstrate that evolutionarily conserved sites are associated with significant decreases in abundance-PCA signal, and that the combination of chemical properties and surface accessibility explains the impact of mutations on folding stability. We then identify activity-biasing mutations using decomposed phospho-PCA scores and show that our data can explain the impact of clinical variants. We further utilize these decomposed scores to identify substrate-specificity-determining residues and to conduct a molecular dynamics study to understand, mechanistically, how mutations at these residues affect substrate recognition. This work represents a significant advance and resource for the field, as it provides a valuable methodology, vast dataset, and furthermore, significant new mechanistic insights. Furthermore, phospho-PCA and the associated computational methods have the potential to generate mechanistic insights for the entire human kinome. Moreover, our comprehensive dataset can support the development and benchmarking of new methods for variant-effect prediction.

## RESULTS

### Phospho-PCA enables deep mutational scanning of kinase domains

Phospho-PCA uses a growth-based biosensor to couple kinase-substrate interactions to the growth of budding yeast cells; a schematic is shown in Figure 1. This biosensor consists of two separate proteins expressed in tandem: the first is a kinase domain tethered to its substrate and one-half of a split dihydrofolate reductase (DHFR) enzyme. The second is a phospho-amino-acid binding domain (PAABD) tethered to the other half of the split-DHFR enzyme^75,76^. When a kinase-substrate interaction occurs, the substrate is phosphorylated, leading to the binding of the PAABD to the substrate. This binding reconstitutes the DHFR enzyme, conferring resistance to methotrexate. We note that this construct design was inspired by fluorescent kinase biosensors, which use a similar mechanism to couple kinase substrate interactions to changes in fluorescence^77,78^. By removing the PAABD, the DHFR fragments complement spontaneously in the cytosol at a rate proportional to the concentration of folded kinase-DHFR1 fusion protein (Figure 1a), thus enabling abundance-PCA measurements^47^. To perform deep mutational scanning, we first cloned a library of kinase variants (made with one-pot saturation mutagenesis) into the kinase portion of our biosensor. Because kinase domains are too long (750-900bp) for short-read sequencing, we stochastically link the library to short-read nucleotide barcodes and perform long-read sequencing to construct a lookup table^79–81^. Short-read sequencing is used to measure the abundance of each variant after methotrexate selection (Figure 1b).

**Figure 1:**
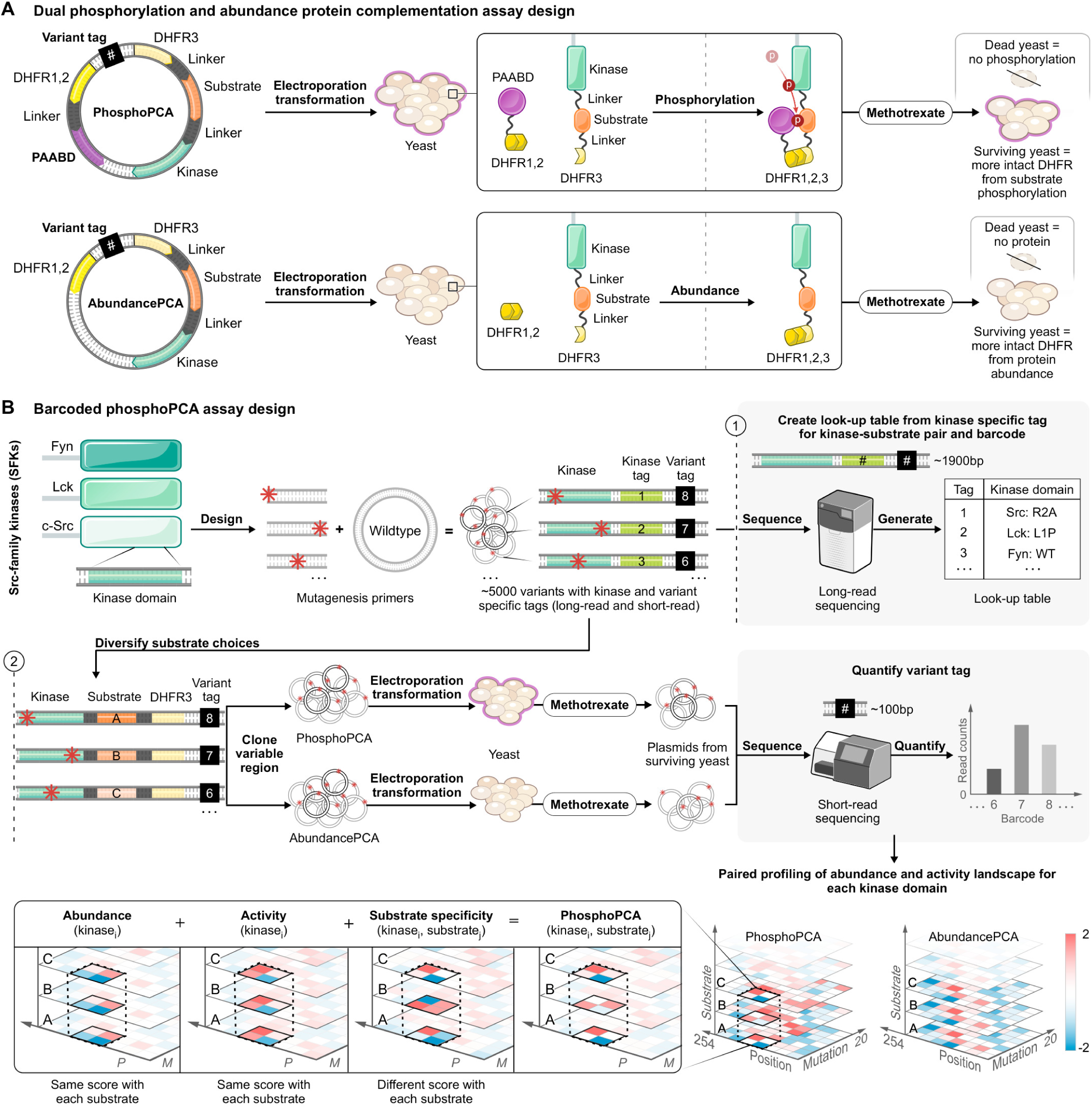
Phospho-PCA enables deep mutational scanning of kinase domains to understand the sequence-to-function relationship for this enzyme category. a) Phospho-PCA utilizes a growth-based biosensor to couple kinase-substrate interactions to cell growth. Kinase-substrate interactions reconstitute the DHFR enzyme, conferring resistance to methotrexate selection. Abundance-PCA can be performed with the same construct by removing the PAABD, providing a score of folding stability. b) Deep mutational scanning of kinase domains with phospho-PCA is performed by cloning a kinase variant library into the biosensor construct and barcoding the variants with 16bp-long barcodes. Long-read sequencing creates a lookup table that links kinase variants to barcodes, where each variant is described by the wild-type amino acid, the position, and the mutated amino acid. Once cloned, the library is transformed into budding yeast cells, and the activity of each variant for phospho and abundance-PCA is measured by quantifying cell growth in the presence of methotrexate with short-read sequencing.

To validate phospho-PCA, we measured growth curves of budding yeast expressing our biosensor construct for a collection of kinase-substrate pairs; the results are shown in Figure 2. We observed significantly higher growth for interacting kinase-substrate pairs than non-interacting pairs (Figure 2a). We used both catalytically dead kinase domains and mismatched substrates to create non-interacting pairs. We validated that our growth-based biosensors worked for the Fyn, Lck, and c-Src tyrosine kinases, a collection of serine-threonine kinases, and for a panel of phospho-amino acid binding domains (PAABDs) (Figure 2a,c). Importantly, by quantitatively extracting growth rates during the exponential phase of cell growth, we observed that our biosensors recapitulated the known preferences of Fyn, Lck, and c-Src for their respective substrates (Figure 2b). While our study focuses on the three SFKs, the data for serine-threonine kinases is a crucial demonstration of phospho-PCA’s scalability. These results demonstrate that we can utilize our growth-based biosensors to quantify the strength of interaction for almost any kinase and substrate during cell growth.

**Figure 2:**
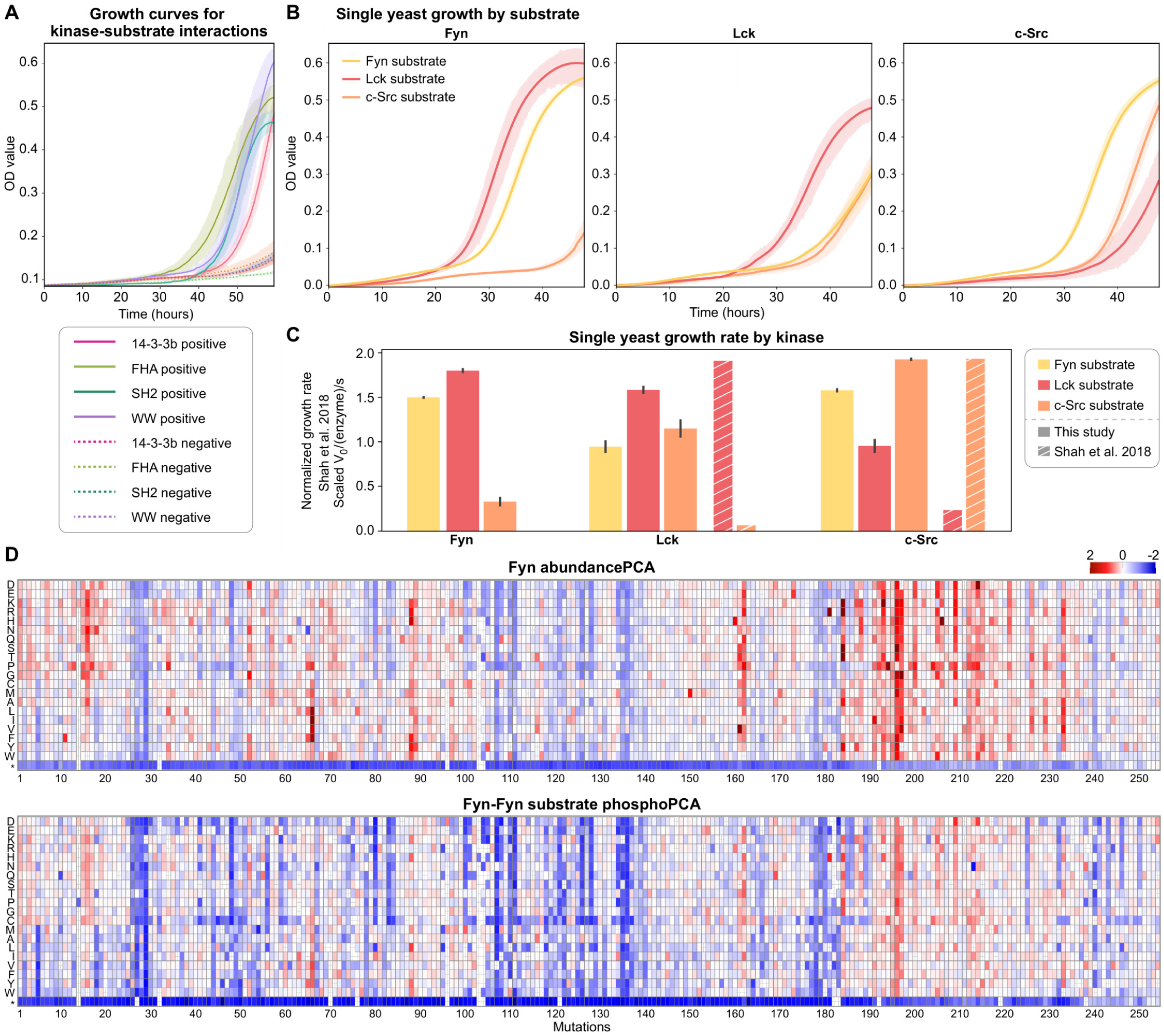
Phospho-PCA quantitatively couples kinase-substrate interactions to cell growth. a) Biosensors containing interacting kinase-substrate pairs resist selection, while biosensors with non-interacting pairs display significantly reduced growth. b, c) Quantitative measurement of Fyn, Lck, and c-Src substrate preferences for Fyn-sub, Lck-sub, and c-Src-sub with phospho-PCA. Growth rates were computationally extracted during the exponential phase of cell growth. Scores for all substrates of each kinase were standardized using z-score normalization and then rescaling to a range of 0 to 2. Kinase-substrate interaction strengths measured with phospho-PCA are consistent with prior measurements of substrate preferences. d) Heatmaps for abundance- and phospho-PCA measurements for Fyn and the Fyn/Fyn-sub interaction, respectively.

With our growth-based biosensors validated, we constructed libraries to explore the impact of variants on activity and substrate recognition for the SFKs Fyn, Lck, and c-Src. For each kinase, we generated a library of all possible single-site variants and paired it with one potential substrate (Fyn-sub, Lck-sub, and c-Src-sub). We constructed 18 biosensor libraries that covered phospho-PCA and abundance-PCA for all possible kinase-substrate pairs. These libraries were screened by measuring growth rates under methotrexate selection using short-read sequencing. In-depth descriptions of our library assembly method, analysis of each library’s mutation composition and coverage, methotrexate selection, protocols, and bioinformatic analysis of the sequencing data are given in the Methods and Supplemental Information.

### Structural context determines the impact of mutations on SFK abundance

To investigate the impact of mutations on SFK folding stability from a biophysical perspective, we performed a principal component analysis of our abundance-PCA data. We note that abundance-PCA captures the impact of variants on folding stability, which reduces the amount of folded kinase inside the cell^47^. For simplicity, we refer to a variants impact on abundance-PCA scores, and hence folding stability, as “abundance” for the rest of this paper. For this analysis, we first grouped each amino acid (excluding proline, cysteine, and glycine) into four biophysical categories: hydrophobic, polar uncharged, positively charged, and negatively charged. We then compiled the biophysical category and abundance data for all mutants across all positions and kinases and performed principal component analysis on the resulting matrix; the results are shown in Figure 3a and 3b. To simplify our description of the results, we adopt the convention of Modi and Dunbrack for the relevant sections of the kinase domain^82^. This convention defines 17 key sections throughout the kinase domain - 8 in the N-terminal domain (B1N, B1C, B2, B3, HC, B4, B5, HD) and 9 in the C-terminal domain (HE, CL - catalytic loop, ALN - activation loop N terminus, ALC -activation loop C terminus, HF, FL, HG, HH, HI). Each section is named for its dominant secondary structure or its functional role. Throughout this work, residue numbers in the text refer to positions within the isolated 254-residue kinase domain construct, which can be mapped to the full sequence by adding the appropriate offset for each kinase (269 for Fyn, 260 for Lck, and 244 for c-Src).

**Figure 3:**
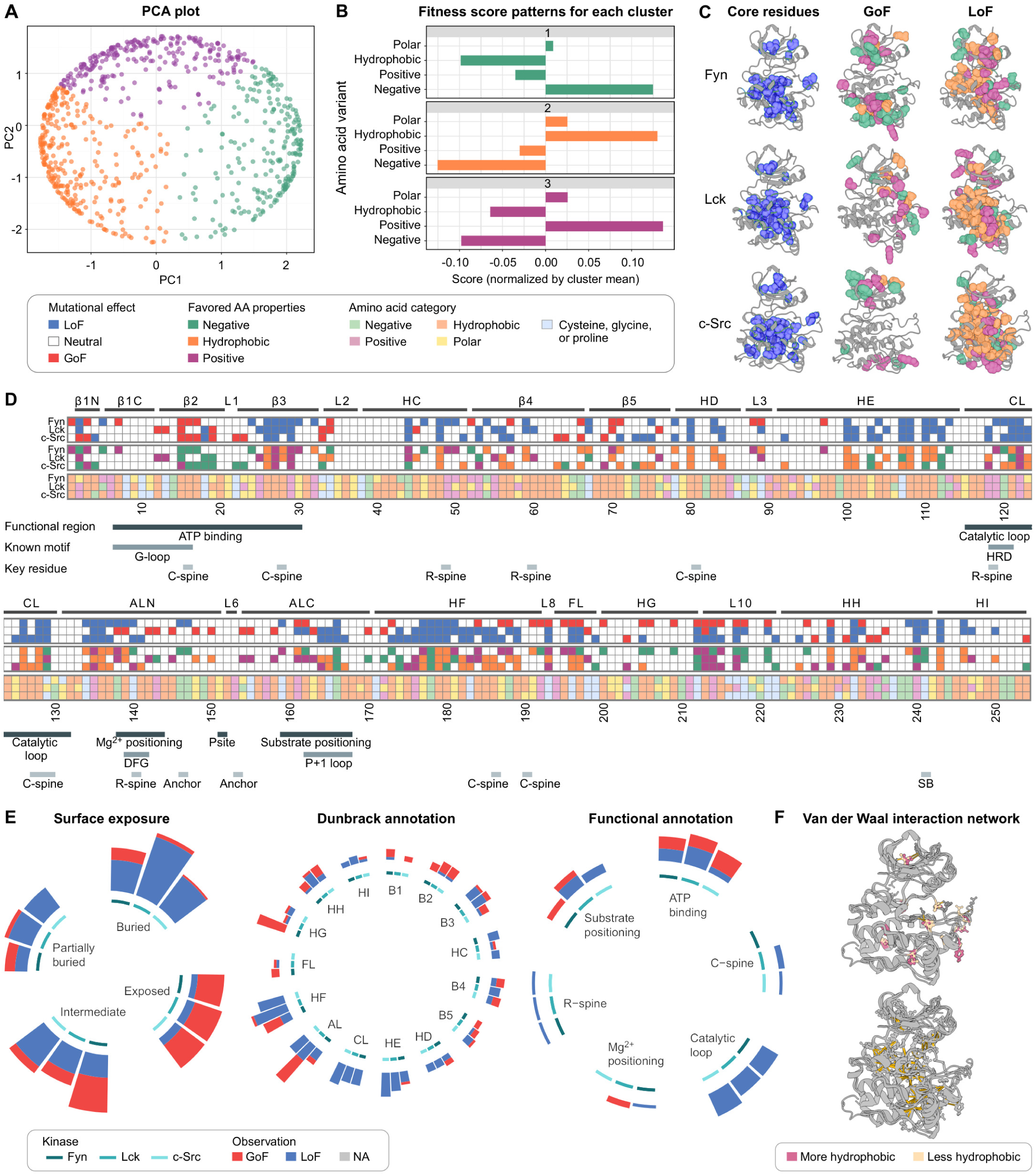
Mutations in conserved regions impact abundance-PCA scores of SFKs. a-b) Principal component analysis reveals three distinct responses of positions to mutations. Group 1 and group 3 positions prefer negatively or positively charged residues, respectively, while group 2 positions prefer hydrophobic residues. c) Positional response categories overlaid on the kinase structure. Group 2 positions cluster at the hydrophobic core, while group 1 and group 3 positions are enriched at the kinase surface. d) Functional annotations and mutational effects for the kinase domains of Fyn, Lck, and c-Src. The six rows display, for each position, the Dunbrack annotation, whether the amino acid property is conserved across the three SFKs, the overall mutational effect, the favored amino acid property, the wild-type amino acid and its category, and a functional annotation. e) Gain-of-function and loss-of-function positions categorized by surface exposure, Dunbrack annotation, and functional annotation. f) Disruption of van der Waals networks explains the impact of mutations on abundance-PCA scores. Top: Kinase-specific van der Waals networks govern solvent accessibility and mutation tolerance in abundance-PCA scores. Bottom: Destabilizing sites conserved across SFKs disrupt van der Waals networks in the hydrophobic core.

Using the principal components, we identified three clusters where the presence of negatively charged (1), hydrophobic (2), or positively charged residues (3) leads to a gain-of-function effect in abundance-PCA, whereas alternative properties result in a loss-of-function effect. We then sought to use this clustering, in conjunction with physical and structural reasoning, to explain the impact of convergent and divergent mutations (i.e., mutations with the same or different effects across all three SFKs, respectively). With this goal in mind, we made several observations. First, we observed that most conserved abundance loss-of-function mutations occurred within the hydrophobic core, although some also occurred in structurally distinct regions, such as the HE region and the catalytic loop. Regions with reduced solvent-exposed surface area (SASA, *<* 10Å^2^) were enriched for abundance loss-of-function mutations. In contrast, regions with more solvent exposure were enriched for gain-of-function mutations (Figure 3c). These observations were consistent with our clustering analysis; cluster 2 sites were located in the hydrophobic core, while cluster 1 and 3 sites were located at the surface (Figure 3c). We also analyzed the impact of mutations on abundance-PCA in functionally important, evolutionarily conserved regions (Figure 3d and 3e). We observed that each functional region had a different mutational tolerance. The catalytic loop, which is important both functionally and structurally, and residues in the N-terminal portion of the activation loop that are responsible for magnesium positioning, were enriched for loss-of-function mutations. In contrast, the region responsible for ATP binding contained both gain-of-function (B2) and loss-of-function (B3) mutations.

We next sought to understand the impact of divergent abundance mutations. We observed significant divergence for abundance gain-of-function mutations across the three SFKs; these sites were enriched in HF-HG for Fyn, AL for Lck, and B2 for c-Src. We also observed that the sites for these divergent mutations overlapped with regions where the electrostatic charge distribution differed considerably between these three kinases (Figure 3d). A closer inspection of these locations revealed a consistent relationship between the physicochemical properties of amino acids and the phenotypic outcomes of mutations. Specifically, at several positions, the introduction of a charged residue that is compatible with the local electrostatic environment of one kinase led to a gain-of-function, while being neutral or detrimental to the other kinases that had a different charge distribution at that site. These sites include positions 172 in cluster 1 (Lck) and 76, 88, 154, and 236 in cluster 3 (Lck and Fyn). Similarly, divergent loss-of-function mutations also reveal location-specific preferences for electrostatic charge. These include mutations in positions 4 in cluster 1 (Src) and 247 in cluster 3 (Lck), suggesting that these kinases evolved to favor negative and positive charges at these positions, respectively.

Finally, we observed that networks of van der Waals interactions were a significant driver of divergence, as they impact the degree of solvent accessibility for specific residues and hence sensitivity to mutations. For example, at position 28, interactions present in Fyn and Lck bury a hydrophobic residue, leading to a loss of function when disrupted. In contrast, c-Src lacks these interactions, leading to increased solvent exposure and greater tolerance to mutation. Identified sites are visualized in 3f; a comprehensive list is provided in Table S2. When examined together, these data present a consistent picture in which the structural context of a residue determines the impact of mutation on SFK abundance. Disruption of residues within the hydrophobic core reduces abundance, whereas mutations that increase the likelihood of favorable solvent interactions increase abundance. These effects can be brought about by directly altering the physicochemical properties of a residue or by indirectly disrupting van der Waals networks that control surface exposure. We note that these observations are consistent with prior work, which demonstrates that solvent interactions can enhance a protein’s thermostability and solubility^83^. Overall, these new findings highlight that the effects of variants on SFK abundance are governed by the local structural environment—particularly solvent accessibility, charge compatibility, and van der Waals networks.

### A hierarchical Bayesian model separates the contribution of kinase abundance, activity, and substrate specificity to phospho-PCA scores

In this section we developed a novel computational method to understand the mechanistic impact of variants on kinase function from our paired phospho and abundance-PCA data. Understanding the mechanism behind a variant’s impact is challenging because there are often multiple pathways a variant can take to perturb a protein’s function. For kinases, we envision three mechanistic paths through which mutations can impact the phospho-PCA score, our measure of kinase function. First, a variant could alter folding stability, thereby impacting its abundance in the cell. This would directly affect both the phospho-PCA score (assuming rate-limited kinetics) and the abundance-PCA score. Second, a variant could drive a dynamic shift towards either an active or an inactive conformation. Third, a mutation could alter a kinase’s substrate specificity. To understand the contributions of each of these possible mechanisms to our measured phospho-PCA scores, we developed a hierarchical Bayesian model. This model, displayed in Figure 3a and described in full detail in the supplement, posits that phospho-PCA scores can be decomposed into a linear sum of scores that capture a variants’s impact on abundance (derived from scaled abundance-PCA measurements), kinase activity (shifts towards active or inactive conformations), and substrate specificity. We reasoned that some of these scores should be shared between conditions. For instance, scores for a variant’s impact on abundance should be shared across substrate identity, as the folding stability of the kinase domain should be independent of which short peptide substrate it is tethered to. We confirmed this expectation with exploratory analysis; we note that this effectively allows us to treat the three substrate-specific abundance measurements as technical replicates of the same underlying abundance parameter. Similarly, kinase activity should also be shared across substrate identity. Scores for substrate specificity, however, are expected to vary among all four experimental variables (kinase, substrate, position, and mutation). This expectation can be viewed as a form of regularization that we incorporated into our model. We fit this model to our data using Markov chain Monte Carlo^84^; an example decomposition is shown in Figure 3b. One advantage of the Bayesian framework for modeling is its ability to quantify uncertainty, which enables the identification of key residues that influence kinase activity and substrate preferences. We note that this decomposition was made possible by the presence of the abundance-PCA data, which was a significant prior advance^85^. Although computational methods for decomposing paired protein complementation assay data exist^86,87^, our need to separate the impact of three distinct mechanisms required a bespoke modeling effort. Heatmaps of decomposed scores for kinase abundance, activity, and substrate specificity are shown in Figure 4b and Figure S10. Our modeling effort is a significant advance, as, to our knowledge, this is the first time an effort has been made to model the relative contributions of variants to abundance, catalytic activity, and substrate specificity for any enzyme.

**Figure 4:**
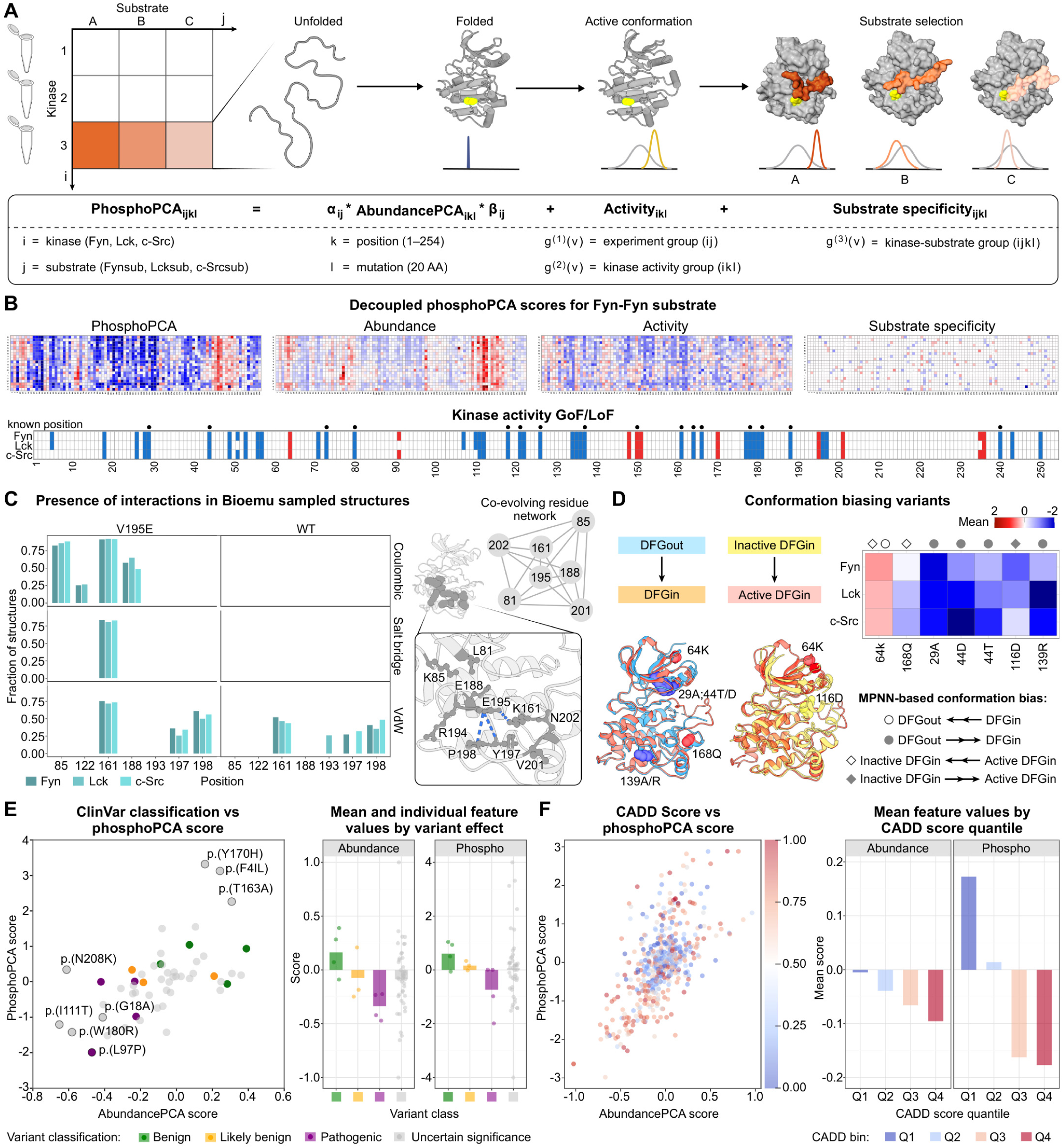
Phospho-PCA reveals activity-biasing mutations and the functional impact of clinical variants. a) Hierarchical Bayesian model schematic for decomposing phospho-PCA scores into abundance, kinase activity, and substrate specificity components. Indices representing kinase identity (i), substrate identity (j), position (k), and mutation (l). b) Example decomposition for Fyn-Fyn substrate pair showing separation into constituent components. Heatmaps display normalized scores revealing mutations that specifically impact kinase activity versus substrate recognition. Variants that impact kinase activity are largely conserved across the three SFKs. c) Conserved gain-of-function mutation V195E reveals a co-evolutionary network. Left: BioEmu conformational sampling reveals interactions among coevolving residues in V195E and wild-type kinases. Right: Network diagram showing V195E establishing new electrostatic interactions that stabilize active conformations. d) ProteinMPNN-based conformational bias analysis identifies mutations favoring different kinase states. Mutations favoring inactive conformations correlate with negative activity scores; those favoring active states correspond to gain-of-function effects. e) Clinical variant analysis using ClinVar. Benign variants cluster near the origin; pathogenic variants show significant negative effects in one or both assays. f) Cross-analysis of CADD scores and experimental results. High CADD variants exhibit reduced abundance and activity, with mean feature values indicating increasingly negative effects as predicted pathogenicity increases.

### Computational decomposition reveals conserved activity-biasing variants

The presence of decomposed kinase activity scores for all three SFKs provides an opportunity to explore how variants impact catalytic activity. Indeed, our analysis of these kinase activity scores identified a set of residues that primarily control SFK catalytic activity, distinct from those essential for abundance. We reasoned that these variants exerted their effect either by directly impacting catalysis or by biasing the kinase toward specific conformations, and used this hypothesis to structure our analysis.

We first sought to categorize residues based on their conserved functional impact across Fyn, Lck, and c-Src. To do so, we examined residues where a loss-of-function (LOF) phenotype was conserved in both abundance-PCA and the decomposed kinase activity scores. We observed that 15 of the 24 conserved LOF positions identified in the abundance screen also showed a conserved LOF impact on kinase activity, suggesting their dual roles in maintaining stability and catalytic function. Of these, ten map to well-known structural motifs and reinforce existing mechanistic insights. For example, V25 and K28 (*β*3) are critical for ATP binding, with K28 forming a key salt bridge with E44 (HC). Other residues with known catalytic roles include E166 (ALC), which is involved in substrate positioning, and R240 (HH), which forms a salt bridge with E166. Our analysis also highlighted residues forming the hydrophobic catalytic spine (C-spine), including L80 (HD), L121, and I126 (Catalytic Loop), whose strong LOF phenotypes underscore the C-spine’s architectural importance. D178 (HF) also showed a strong LOF phenotype, consistent with its role in stabilizing the strained backbone conformation of the HRD motif through hydrogen-bonding interactions with the backbone amides of H145 and R146^88^, a conserved feature critical for maintaining the catalytic loop geometry required for kinase activity. Beyond these motifs, our screen also identified five previously uncharacterized but essential residues, such as I/V136 (ALN) which is adjacent to the R-spine in the Mg^2^+ positioning region of the activation loop, a cluster on the HF helix (V179, W180), and F243 on the HI helix, suggesting they have unappreciated roles in kinase structure and regulation.

To identify residues whose influence is confined to catalytic regulation, we focused on positions exhibiting a conserved LOF phenotype exclusively in the decomposed kinase activity scores. These residues suggest a specialized role in modulating kinase activity or conformation rather than contributing to overall protein folding. We identified a total of 14 residues, 8 of which have known roles in catalysis. These include G18 (*β*2 strand), I28 (*β*3 strand), and E73 (*β*5 strand) for ATP binding; A137 (xDFG motif) for Mg^2+^ coordination; and K161 and A164 (ALC) for substrate alignment in the P+1 loop. Notably, the HF helix, which our analysis identified as enriched for conserved variants, contains critical residues, such as E188 and S181. In PKA, the corresponding mutant to E188 (E230Q) crystallizes in an inactive state even with ATP present, highlighting its role in stabilizing ATP binding^89^. Similarly, S181 on the HF helix helps position the catalytic loop for efficient transfer of the phosphate group. Our screen also revealed six previously unannotated residues with activity-specific LOF effects. L56 and V57, located near the *α*C–*β*4 loop, likely stabilize the R-spine assembly, a key interface for maintaining an active conformation^30^. Other newly implicated residues, including V71 (adjacent to gatekeeper), E112, Y197, and L250, occupy helices in the C-lobe that shape the active-site cleft, suggesting they are important for the dynamic motions required for activation.

While LOF mutations were common, we also identified conserved gain-of-function (GOF) residues that offer insights into activity regulation. For example, at position 150 (Y419 in c-Src), the role of phosphorylation in regulating catalytic activity is more nuanced than simple on/off switching. While Y419 phosphorylation is conventionally associated with activation, non-phosphorylatable mutations retain substantial catalytic activity in certain contexts and can even reduce competition with substrate phosphorylation in single-domain in vitro assays^90,91^. However, Y419 phosphorylation is critical for higher-order functions, particularly dimerization and assembly of macromolecular complexes that modulate substrate access and selection in vivo^91^, explaining why mutations at this site have context-dependent effects on cellular signaling. At position Y170, a known autophosphorylation site, the phospho-mimetic mutation Y170E was enriched with a positive phospho-only score. This suggests that introducing a negative charge may induce the kinase to adopt an active conformation. Novel GOF sites were also identified at 64K and V195E. Interestingly, V195E appeared to stabilize the active state by forming a strong Coulombic interaction and a new hydrogen bond with K161 in the P+1 loop. This interaction functionally offsets the positive charge of K161, a mechanism previously shown to increase Src activity^92^. We explore both of these novel variants in the following section.

Finally, our analysis of canonical catalytic motifs such as HRD, DFG, and APE revealed that they were enriched with mutations that had a mild effect on abundance but a substantially negative impact on the phospho-PCA score^93^. This resulted in negative phospho-only scores for mutations like p.(F139R), p.(E166N), and p.(P165H), confirming their critical and direct role in phosphorylation. In contrast, mutations in other motif residues, such as p. (R119P) and p. (D120P), had strongly negative scores in both assays, indicating a mixed effect on catalysis and abundance. In total, these data highlight the ability of our method to separate variants that purely disrupt catalysis from those that also compromise abundance.

### AI-enabled structural analysis reveals conformation-biasing variants

To understand the structural basis of activity-biasing mutations, we applied recently developed computational methods for conformation stability analysis. These methods utilize the inverse folding model ProteinMPNN to predict a protein’s preference for specific conformations^94,95^. Here, we utilized Protein-MPNN to investigate whether a specific mutation would introduce a bias towards a particular kinase conformation. For this analysis, we used available crystal structures of c-Src and Lck kinases in active (Src:1YI6, Lck:3LCK), inactive DFG-in (Src:2SRC, Lck:3B2W), and inactive DFG-out conformations (Src:2OIQ, Lck:6PDJ). For each possible single amino acid substitution, we computed the preference for inactive DFG-in versus inactive DFG-out conformations, as well as active DFG-in versus inactive DFG-in conformations. This analysis revealed a set of eight variants that are predicted to favor specific conformational states across both kinases; all have a significant effect on activity in our decomposed data. Of these eight, variants 29A, 44D/T, and 139A/R were predicted to favor the inactive DFG-out conformation over the active DFG-in conformation, consistent with their negative kinase activity scores in our experimental data. Conversely, variant 116D was predicted to favor the inactive DFG-in state over the active DFG-in state, also correlating with reduced activity scores. Notably, variants 168Q and 64K were both predicted to strongly favor the active DFG-in conformation over the inactive DFG-out state. However, only 64K was additionally predicted to favor the active DFG-in conformation over the inactive DFG-in state. This prediction aligns with our experimental observation that 64K represents a novel gain-of-function mutation across all three SFKs, while 168Q shows mild loss-of-function effects. We note that although all eight variants identified by ProteinMPNN were present in our data, it failed to identify most of the allosteric sites that impact kinase activity discovered in our study. This indicates ProteinMPNN has a low recall and high precision for variant effect prediction in this setting. We performed a similar analysis of protein language models’ ability to detect allosteric sites and observed results similar to those seen for ProteinMPNN; this analysis is described in more detail in the supplement. While the precision of these methods enables them to provide useful insights into how variants behave, their limited performance suggests that significantly more data are necessary before they can be used more broadly to investigate the impact of allosteric variants.

We also investigated the molecular basis of the V195E gain-of-function mutation using evolutionary coupling analysis and equilibrium conformation sampling (Figure 4c,d). Evolutionary coupling analysis applied to Fyn, Lck, and c-Src revealed a co-evolutionary network involving residues 81, 85, 161, 188, 195, 201, and 202 (Figure 4c, right). To understand how the V195E mutation affects this network, we used BioEmu^96^, a recent method for accelerated generation of conformational ensembles for medium-sized proteins, to sample 4,000 conformations for each wild-type kinase and their corresponding 195E mutants. Structural analysis of the generated ensembles revealed that the 195E mutation establishes new Coulombic interactions with residues 85, 122, 161, and 188. Most significantly, a salt bridge between residues 161 and 195 was prevalent only in mutant conformations, bringing these residues into close proximity (Figure 4c, left). To explore the impact on kinase activity, we used Kincore to classify conformations as active or inactive. We then evaluated the percentage of structures that were classified as active, depending on whether the salt bridge was present or absent. This percentage for conformations with and without the salt bridge were: 8.1% and 2.9% for Fyn, 2.7% and 0% for Lck, and 6.1% and 0% for c-Src, respectively. These results demonstrate that the V195E mutation enhances kinase activity by stabilizing active conformations through the formation of a critical salt bridge within a network of co-evolving residues.

### Abundance and phospho-PCA scores reveal the impact of clinical variants

A wide range of human diseases can arise from single-point mutations in SFKs, and for many human variants, the potential pathogenicity and function are unknown. We therefore reasoned that integrating abundance and phospho-PCA scores could provide a powerful framework to connect human kinase variants to their molecular and functional consequences. To do so, we first analyzed the behavior of known benign and pathogenic variants across the three SFKs from the ClinVar database within our experimental data (Figure 4e)^97^. Benign variants displayed near-zero to mildly positive abundance and phospho-PCA scores, consistent with preserved protein function. In contrast, pathogenic variants frequently displayed significantly negative values in one or both metrics, underscoring the essential role of SFKs in human biology. Several variants exhibited negative abundance scores, accompanied by only slight changes in phospho-PCA scores, suggesting that reduced protein stability alone can drive pathogenicity. We speculate that the non-catalytic function of SFKs may explain this observation. Furthermore, several pathogenic variants exhibited significantly negative scores for both abundance- and phospho-PCA. Interestingly, these trends can be used to infer the pathogenicity for variants of unknown significance (VUS). For instance, the VUS variant N208K displayed a pronounced reduction in abundance-PCA while the VUS variants I111T, W180R, and G18A display significant reductions in both abundance and phospho-PCA, suggesting that all four are pathogenic variants.

To expand our analysis beyond Clinvar’s SFK variant collection, we collected population allele frequency data from the Genome Aggregation database (gnomAD)^98^ and annotated them with pre-computed Combined Annotation Dependent Depletion (CADD) scores^99^, a computational prediction of how harmful a genetic variant is likely to be. We then used these CADD scores to continue our analysis of the functional impact of SFK variants (Figure 4e). When overlaid on our experimental data, where we plot the abundance and phospho-PCA scores for each variant, we observed that variants with low CADD scores (presumably benign) clustered near the origin. However, high CADD variants were concentrated in the bottom-left quadrant, with negative scores for both assays. This finding was consistent with our analysis of the ClinVar database, which revealed a correlation between variant pathogenicity and a negative impact on both scores. Interestingly, the top-left quadrant contained a smaller number of high-CADD variants with reduced abundance-PCA scores but high phospho-PCA scores. This suggests that these rare variants may be deleterious gain-of-function mutations. This analysis demonstrates that the combined abundance and phosphorPCA scores can provide crucial insights into the function of clinically significant variants.

### Molecular dynamics reveals allosteric networks perturbed by specificity determining variants

Allosteric effects, where perturbations at one site propagate through a protein to affect distant functional regions, are ubiquitous in biology; understanding how allosteric communication unfolds and is perturbed by variants is critical for understanding protein evolution and disease. Our computational decomposition of phospho-PCA generated scores for substrate specificity, providing an opportunity to explore the role of allostery on substrate preference. We identified a collection of variants that specifically impact substrate preference for these three SFKs; these residues are shown in Figure 5a and described in full detail in the supplement (Table S3). From the statistically significant variants we identified, we performed additional filtering by categorizing amino acids based on their chemical properties. We then identified variants in which the amino acid property matched one kinase’s wild-type property at that site, but differed from the other two; this yielded a list of variants that induced a clear, directional change in the preferred substrate. Surprisingly, when displayed on top of the protein structure, most of the identified residues were distal to the expected binding interface of the substrate (Figure 5a, right). Based on this observation, we concluded that allostery is likely a key mechanism underlying how these variants affect substrate specificity.

**Figure 5:**
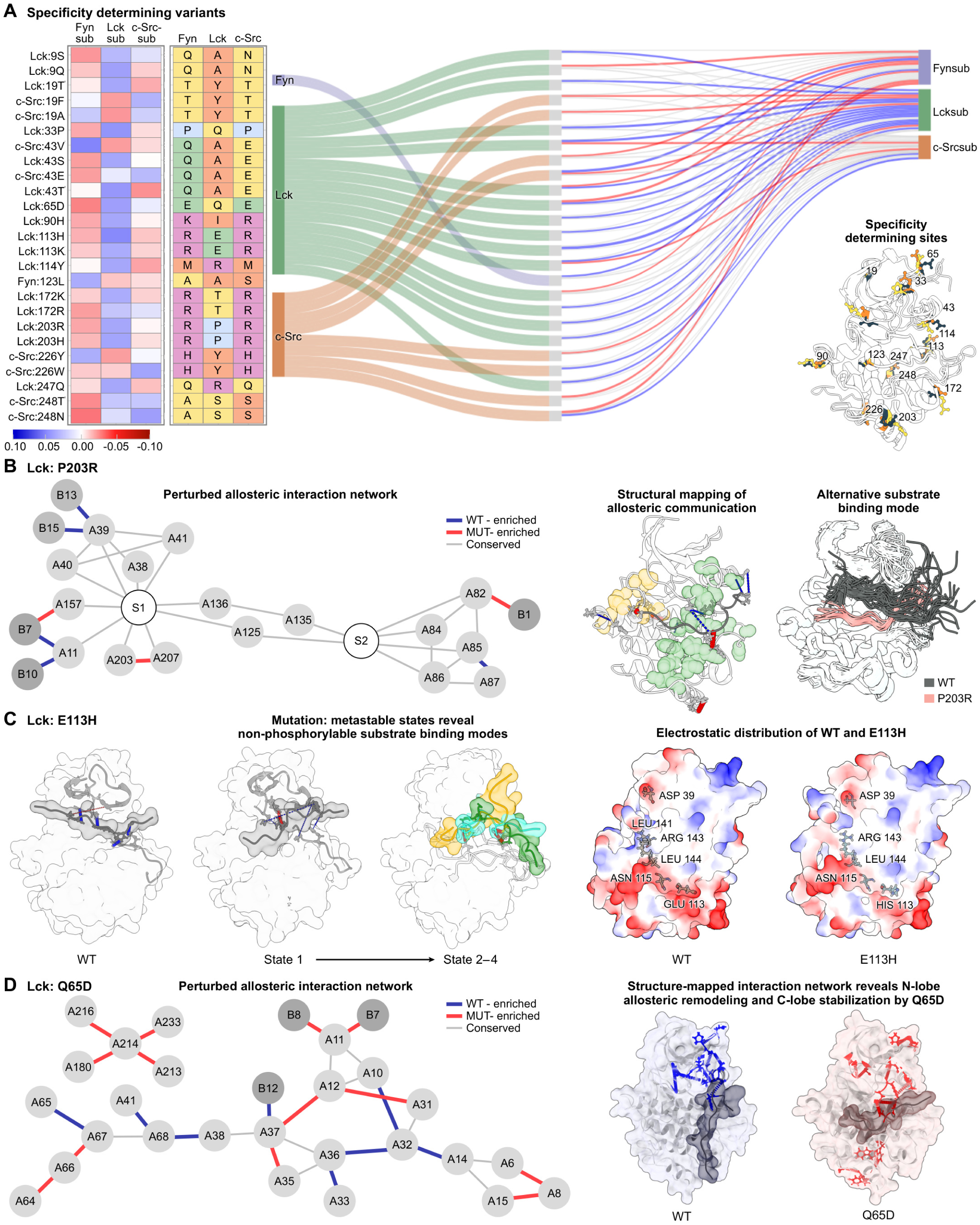
Molecular dynamics reveals allosteric networks governing substrate specificity. a) Specificity-determining variants identified through computational decomposition. Left: Heatmaps showing residue chemical properties and substrate specificity scores. Middle: Sankey diagram connecting specificity-determining mutations to affected allosteric sites. Right: Structural mapping showing most specificity-determining residues are distal to the kinase-substrate interface. b) Lck P203R rewires an allosteric network, which leads to altered substrate positioning. Left: A residue interaction graph shows communication pathways from HG through the HF-HG loop → P+1 loop → catalytic/activation loops → N-lobe. Kinase residues are labeled A and substrate residues are labeled B. S1 and S2 are supernodes connecting correlated residues for visualization. Middle/Right: Structural mapping and frame overlay show that the substrate shifts toward the C-lobe, with the phosphorylatable tyrosine farther from ATP. c) Lck E113H enables an electrostatic switch reshaping binding modes. Left: Overlay of WT and four E113H macrostates shows phosphorylation-incompetent poses. Right: Electrostatic surface comparison shows the E → H substitution creates a patch of altered electrostatic potential along the A113–A115–A144–A143/141 pathway, destabilizing productive substrate positioning. d) Lck Q65D re-configures N-lobe packing to favor Fyn-like substrate. Left: A residue interaction network showing differential interactions between Q65D and wild-type. Right: Structure comparison reveals N-lobe allosteric remodeling and C-lobe stabilization from introduced negative charge at *β*4–*β*5 turn.

To understand how distant mutations alter substrate specificity through allosteric mechanisms, we performed molecular dynamics (MD) simulations of several specificity-determining variants bound to their substrates. We used the Chai-1 protein folding model to generate kinase-substrate complexes and simulated both wild-type and variant kinases for 500 ns using GROMACS^100,101^. Our analysis focused on Lck and c-Src, as we were unable to obtain confident structural models for Fyn in complex with its substrate. Details of the structure generation and MD simulation protocols are provided in the Methods section. Our central goal was to trace how each distant specificity-determining mutation communicates with the substrate-binding interface—that is, to map the allosteric pathways connecting the mutation site to the residues that directly contact the substrate. To accomplish this, we developed a contact graph framework that identifies plausible routes of structural communication through the protein. First, we constructed residue contact graphs for each simulation, where nodes represent individual residues and edges connect residue pairs that form frequent contacts during the trajectory^102^. We then identified contacts that were strengthened (red edges) or weakened (blue edges) in the variant relative to wild-type, highlighting how the mutation rewires the kinase’s internal interaction network. Next, we defined a set of “critical residues” that must be connected by any viable allosteric pathway: (1) the mutated residue itself, (2) substrate residues whose kinase contacts differ between variant and wild-type, (3) kinase residues that directly contact the substrate, and (4) residues in structural regions showing significant changes in correlated motion between variant and wild-type simulations. Finally, we identified minimal pathways connecting all critical residues by solving an approximation to the Steiner tree problem^103^, which finds the smallest subgraph linking a specified set of nodes. These pathways reveal how structural perturbations propagate from the mutation site through intermediate contacts to ultimately reshape substrate recognition at the binding interface. A detailed description of the contact graph construction, critical residue identification, and pathway extraction is provided in the Methods.

The results of this analysis are shown in Figure 5 and revealed distinct mechanisms by which different specificity-determining variants use allosteric pathways to modulate substrate preference. Some variants induced new substrate-binding modes by reorganizing interface contacts, while others introduced new inactive metastable conformations that reduced overall catalytic competence. We describe the specific mechanism for each analyzed variant below.

- *Lck P203R introduces a new, inactive binding mode for the Lck substrate.* Position 203 in Lck represents a critical allosteric site controlling substrate specificity through long-range communication networks within the kinase domain (Figure 5b). The P203R mutation reduces Lck’s affinity for its native substrate while enhancing activity toward Fyn/c-Src substrates, consistent with prior work showing that the corresponding c-Src residue (R472) coordinates substrates containing negative charges downstream of the phosphorylation site^22^. Our molecular dynamics simulations revealed that P203R dramatically repositions the substrate closer to the C-lobe and farther from ATP, accompanied by specific interaction changes at positions +1, +7, +10, +13, and +15—the same positions differing in charge distribution between Lck and c-Src substrates. The mutation propagates its effects through a sophisticated allosteric network: the arginine substitution establishes new intramolecular interactions (sidechain hydrogen bonding between residues 203-207 and increased 205-197 contacts), which alter interaction frequencies along the 203-205-197-196-161-159 pathway, transmitting from HG through the flexible HF-HG loop allosteric hub to the P+1 loop and ultimately to N-terminal lobe through regulatory spine interactions (e.g., via HRD/DFG R-spine residues).
- *Lck E113H creates new, inactive metastable states.* Position 113, located five residues upstream of the HRD motif in the catalytic loop, exhibits divergent electrostatic properties among SFKs (glutamic acid in Lck versus arginine in Fyn/c-Src) and has been previously implicated in controlling substrate specificity through charge-dependent peptide recognition^22^. Our molecular dynamics simulations revealed that the E113H charge reversal fundamentally alters Lck’s tolerance for positively charged substrates by stabilizing non-productive binding conformations (Figure 5c). Across two independent simulations, E113H consistently adopted four distinct phosphorylation-incompetent metastable macrostates with altered substrate positioning compared to wild-type, disrupting critical N-terminal activation loop contacts (Arg143-substrate +12 and Leu141-substrate +11) that normally position the substrate optimally for phosphoryl transfer. Residue interaction graph analysis revealed that the perturbation propagates through the 113-115-144-143/141 pathway, affecting both the activation loop’s positioning and HC organization, which, in turn, reduced contacts between HC residues 40/41 and G-loop residue 12. These conformational changes cause looser substrate docking and formation of alternative salt bridges (Arg143-substrate Asp+9 in metastates 2 and 4; Asp120-substrate Arg+5 in metastates 3 and 4) that function as kinetic traps, sequestering the substrate in catalytically incompetent conformations and explaining the experimentally observed reduced phosphorylation efficiency.
- *Lck Q65D dynamically sculpts the substrate-binding interface through allostery.* The Lck Q65D variant exemplifies a specificity switch, causing loss of preference for the Lck substrate while enhancing activity toward the Fyn substrate through both altered surface electrostatics and allosteric network communication (Figure 5d). The Q65D substitution at the *β*4-*β*5 turn creates a new negative charge on the N-lobe surface near the substrate entrance that electrostatically attracts the multiple positive charges upstream of the phosphorylatable tyrosine in the Fyn substrate, enhancing encounter rates and binding affinity. Additionally, Q65D modulates substrate selectivity by altering the kinase’s preference at the -1 position from bulky hydrophobic residues (Lck substrate’s leucine) to polar residues (Fyn substrate’s threonine) through allosteric rewiring of HC/*β*-sheet coupling networks. This rewiring involves a cascade of N-lobe interaction changes (reduced A65-A67 contacts, gained A64-A66-A67 interactions, disruption of the A68-A41-A38-A14-A32-A33-A36 hydrophobic network) that reposition residue A11 to accommodate polar residues at position -1 through enhanced van der Waals interactions. The improved substrate anchoring is further stabilized by new C-lobe networks involving A213, A214, and A233, with closer contacts to A180—a residue in the HF segment that is part of the tyrosine kinase hallmark sequence (DVWSFG) critical for P+1 pocket geometry. These C-lobe contact changes emerge concomitantly with the N-lobe *β*4-*β*5 rewiring, where the altered electrostatic landscape and loop conformations collectively reshape the peptide-binding pose. This coordinated remodeling of the N-lobe and C-lobe suggests that the divergent HG-HH region plays a key role in determining kinase-specific substrate preferences.

Collectively, these three variants exemplify how diverse allosteric mechanisms—spanning electrostatic remodeling, metastable-state stabilization, and network rewiring—enable them to exert their influence from distal sites to reshape the substrate-binding interface.

## DISCUSSION

This study represents the most comprehensive experimental characterization of how mutations impact folding stability, catalytic activity, and substrate specificity for any enzyme family to date. We systematically measured the functional impact of over 15,000 kinase domain variants by profiling more than 45,000 kinase-substrate interactions across nine kinase-substrate combinations (three kinases paired with three substrates each), generating phospho-PCA and abundance-PCA measurements for each. This scale is matched by the depth of our mechanistic insights, as our approach disentangles the contributions of folding stability, catalytic activity, and substrate preference for each variant through hierarchical Bayesian modeling. By exploring the sequence-function relationship for three SFKs and their variants, we gained mechanistic understanding of how variants alter kinase conformational ensembles and how allostery shapes the kinase-substrate interface. Critically, we found that exploring conformation landscapes through computational simulations was essential for rationalizing our observations. These simulations, which included conformational ensemble sampling and molecular dynamics, revealed sophisticated allosteric communication networks that connect distal variants to changes in activity and substrate specificity.

Beyond its scale, our work has important implications for protein biology, variant-to-function studies, and computational modeling of allostery. The insights gained from this study on protein kinases are also important for other enzyme families, as the principles governing how structural context determines mutational impact on folding stability, and how allosteric networks propagate functional changes, are likely universal. Our comprehensive dataset provides a useful resource for the community, enabling bench-marking of variant-effect prediction methods and providing ground-truth data for understanding the molecular principles governing enzyme specificity. Indeed, evaluation of existing AI methods, including protein language models and inverse folding approaches, for identifying allosteric sites revealed limited predictive power (Figures 4d and S3), as the methods we explored had high precision but low recall. This highlights a crucial limitation: the evolutionary information that self-supervised models learn to extract from protein sequences can differ substantially from the functional properties of interest. Our work addresses this need by demonstrating a scalable approach to generating high-quality functional data on protein variants that goes beyond measures of activity. Combining such functional and biophysical information with protein language models may offer a promising path forward for improving computational predictions^104^. Moreover, our biosensor design, which uses a physical interaction between a modified substrate and a sensor domain to link enzyme activity to cell growth, may be generalized to other enzymes responsible for post-translational modifications^105^, providing a scalable framework for mapping functional landscapes across entire enzyme families.

In addition to advancing fundamental understanding of enzyme function, our work provides actionable insights for clinical variant interpretation and drug discovery. Understanding the variant-to-function relationship is essential for interpreting genetic variation in human populations and understanding the genetic basis of human disease. Our analysis of ClinVar and gnomAD data strongly suggests our comprehensive functional map can be used to interpret clinical variants, as we observed clear signatures that distinguished benigh from pathogenic variants. Beyond variant interpretation, our insights into the role of allostery in protein kinases will inform therapeutic development. Traditional kinase inhibitors target the ATP-binding pocket, often leading to off-target effects due to the structural similarity of kinase domains across the human kinome^106^. Our identification of allosteric sites linked to catalytic activity may provide a path to more selective kinase inhibition^107–109^. Moreover, our identification of distal allosteric sites that modulate substrate specificity without broadly affecting catalytic activity suggests a fundamentally different strategy for drug design: developing highly selective kinase modulators that alter substrate preference rather than simply blocking activity. Such allosteric modulators could enable more precise therapeutic interventions by selectively disrupting pathogenic signaling while preserving beneficial kinase functions, potentially reducing the side effects that have limited the clinical utility of many kinase inhibitors. The allosteric mechanisms we uncovered—including electrostatic switches, conformational biasing through dynamic network rewiring, and metastable-state stabilization—may provide specific molecular handles for the rational design of next-generation kinase therapeutics.

We close by noting that our work demonstrates that phospho-PCA can be scaled to the entire human kinome. The growth-based biosensor works for both major kinase families, and the method requires only knowledge of kinase-substrate interactions to perform deep mutational scanning. We estimate that each barcoded kinase library costs ≈ 3000 USD in long-read sequencing costs, and each mutational scan costs an additional ≈ 1000 USD in short-read sequencing costs. Extending measurements of a similar scale (phospho-PCA measurements for three substrates and one abundance-PCA measurement per kinase) to the entire human kinome would cost ≈ 3.5 × 10^6^ USD. Although the cost is substantial, it is reasonable considering the value of the data. Such a dataset would provide insights into information transmission in signaling pathways, link variants of unknown function to human disease, provide broader insights into allosteric network organization, and facilitate the development of novel therapies targeting allostery. Such a comprehensive functional map would significantly advance our understanding of how sequence variation shapes cellular signaling networks; the work presented here now places such an advance within reach.

### Limitations of the study

There are several limitations of phospho-PCA and our study that merit discussion. First, we leverage yeast as the model organism for phospho-PCA. The cellular environment (e.g., pH, ionic strength, and cofactor concentrations) in yeast differs from that in mammalian cells, which could impact the translation of our results. Further, yeast express a number of endogenous serine-threonine kinases; this background signal may pose a challenge for future phospho-PCA measurements of serine/threonine kinases. We note that our Bayesian approach to separating the impact of abundance from phospho-PCA scores also has limitations. This model assumes that phospho-PCA scores can be decomposed linearly into abundance, activity, and specificity components, which may oversimplify the relationships between these factors. Another limitation is that the difference in scale between activity and specificity scores suggests that phospho-PCA is more sensitive to changes in catalytic rates than to changes in substrate binding affinities. This is likely due to the tethered design of the phospho-PCA biosensor, which creates a high local substrate concentration. While our hierarchical Bayesian framework can still reliably detect mutations that substantially alter substrate specificity, this scale difference prevents conversion of decoupled scores into effective differences in free energy. Our analysis of clinical variants relies on a connection between the functional scores measured in yeast and human disease phenotypes. Future work should seek to further validate this connection, as this link likely depends on cellular and tissue contexts not captured by our system. While phospho-PCA can identify variants with allosteric effects, it cannot directly map the long-range cooperative networks that mediate allostery. While we leverage molecular dynamics to infer these networks, this approach also has limitations in terms of accuracy and scalability. Our study focused on the kinase domain itself and did not examine the effects of mutations in non-catalytic domains, which play a key role in kinase regulation. Indeed, prior work has shown that kinase-domain variants can alter interactions between the kinase domain and other domains^55^. Although our method can, in principle, be extended to perform deep mutational scans on the full-length kinase, we do not explore this here and leave it to future work. Lastly, our emphasis on single amino acid variants means that our study cannot address epistatic interactions between residues. We also leave this to future work.

## Acknowledgments

We thank Todd Peterson for the conversations that sparked the idea for this project. We thank Paul Sternberg, Vincent Tagliabracci, Benjamin Neal, Michael Baym, Uri Manor, and Allyson Evans for helpful discussions and critical feedback. David Van Valen thanks Tiny Nanazian, Della Van Valen, Fatemeh Ranjbar, and Bahar Behsaz for inspiration and support during this project. We thank Bricelyn Strauch for her help with illustrations and figures.

## Author contributions

CY and DVV conceived the project. CY, EP, and DVV designed the experiments. CY performed the experiments. CY and DVV designed the sequencing analysis and the method for computationally decomposing phospho-PCA mutational data. CY implemented the data analysis code. CY and DVV analyzed the data. CY, EP, and DVV wrote the paper. DVV supervised the project.

## Use of artificial intelligence

The authors made use of generative AI tools during the preparation of this manuscript, including Claude, Gemini, ChatGPT, and Refine.ink. These tools were used to assist with copy editing, refactoring text, and evaluating manuscript quality. After using these, the authors reviewed and edited the content as needed and take full responsibility for the content of the published article.

## Data and materials availability

Plasmids for the growth-based biosensors described in this work have been deposited at Addgene and are available for non-commercial use. The sequencing reads for the experiments described in this work are available at https://doi.org/10.5281/zenodo.17541172 (short-read sequencing) and https://doi.org/10.5281/zenodo.17541166 (long-read sequencing); the processed data can be found at https://doi.org/10.5281/zenodo.17539481 (analysis pipelines and processed datasets). Movies of our molecular dynamics simulations can be found at https://doi.org/10.5281/zenodo.17539481. The code for our bioinformatics pipelines, which include sequencing analysis, computational decomposition of abundance and phospho-PCA data, and figure recreation, can be found at https://github.com/vanvalenlab/sfk_dms_paper.

## METHODS

### Method details

#### Yeast strain

All experiments were conducted using *Saccharomyces cerevisiae* strain BY4742 (MAT*α* his3Δ1 leu2Δ0 lys2Δ0 ura3Δ0)

#### Single yeast plasmid construction

Plasmid construction began with two protein complementation assay (PCA) backbones: pGJJ001 (binding-PCA containing DHFR fragments under dual CYC promoters) and pGJJ045 (abundance-PCA containing DHFR fragments under CYC and GPD promoters, convertible into phospho-PCA backbone by phospho-amino acid binding domain insertion). An SH2 domain was first introduced into each backbone using BamHI/SpeI restriction sites, generating pCY001 (CYC/CYC configuration) and pCY002 (GPD/CYC configuration). Active and catalytically inactive Fyn kinase domains were subsequently cloned via NheI/HindIII sites, yielding pCY003–004 from the pCY001 backbone and pCY005–006 from the pCY002 backbone. Using pCY005 as the template, a comprehensive panel of kinase–substrate fusion constructs (with a serine-glycine linker connecting the two) was assembled through an identical NheI/HindIII cloning strategy. This approach generated plasmids pKMS003–008 and pKMS012–016, encompassing all combinations of Fyn, Src, and Lck catalytic domains paired with their respective substrate sequences (exemplified by fyn-gs-srcsub, src-gs-lcksub, lck-gs-fynsub). Control plasmids containing catalytically dead kinase domains (pCY006, pKMS015/016) were constructed in parallel to establish baseline measurements.

#### Yeast transformation

Growth-based biosensors were introduced into yeast through transformation with the corresponding plasmid. Yeast cultures were inoculated in 6mL YPD medium and grown overnight at 30°C. Cultures were diluted to OD_600_ = 0.1 in 180mL YPD and grown to OD_600_ = 0.5–0.6. For lithium acetate/polyethylene glycol transformation, approximately 45 mL of cells were harvested, washed twice in 100 mM lithium acetate, and aliquoted into 1.5 mL microcentrifuge tubes. Salmon sperm DNA was prepared by boiling for 5 minutes, followed by incubation on ice. Transformation mixtures were prepared by sequential addition to cell pellets: 240 µL 50% PEG-3350, 36 µL 1 M lithium acetate, 10 µL salmon sperm DNA (10 mg/mL), and 74 µL plasmid DNA (1–5 µg in a total volume of 360 µL). After vortexing, samples were incubated at 30°C for 30 minutes, heat-shocked at 42°C for 30 minutes, pelleted, resuspended in 150 µL water, and plated on SC-URA selective medium for 2–4 days.

#### Quantitative growth-based kinase-substrate interaction assays

Transformed colonies were picked into 96-deep-well plates containing 500 µL SC-URA/MET/ADE medium per well and grown overnight at 30°C. Cell density was measured by transferring 100 µL culture plus 100 µL fresh medium to flat-bottom 96-well plates. The remaining cultures were preserved as glycerol stocks (30% final glycerol concentration) and diluted to OD_600_ = 0.1. For kinase activity measurements, 5 µL diluted cells were added to 95 µL competition medium (1.053× SC-URA/MET/ADE supplemented with 200 µg/mL methotrexate) to achieve a starting density of OD_600_ = 0.005. Growth was monitored for 60 hours at 30°C using a Tecan plate reader with OD_600_ measurements every 15 minutes.

Time-course data were smoothed using a centered moving average (25-point window). Following exclusion of an initial burn-in period (100 smoothed points), exponential growth phases were identified by computing the numerical derivative of smoothed optical density with respect to time. Time points exceeding a dynamic threshold (*τ*, initially 5 × 10^−3^, iteratively reduced by a factor of 1.25) were selected until at least 25 points were retained. When insufficient points met the criteria or the final OD_600_ was below 0.05, the entire post-burn-in trace was analyzed. Growth rates were calculated as the slope across selected exponential-phase windows. To allow cross-kinase comparisons, growth rates were normalized by the z-score within each kinase and then scaled to the range [0, 2]. Condition-level summaries represent median normalized replicate values with variability expressed as standard deviation unless otherwise specified.

#### Construction of deep mutational scanning libraries

To create barcoded biosensor libraries, we first constructed modular screening plasmids from the abundance-PCA and phospho-PCA plasmids by introducing a BamHI site upstream of the (GGSGG)_4_ linker using KpnI and NheI restriction sites. These modifications generated matched kinase-substrate PCA constructs: pDMS001 (abundance-PCA; Fyn kinase with Fyn substrate) and pDMS002-004 (phospho-PCA; Fyn, Lck, and Src kinases with their respective substrates). Kinase open reading frames were subcloned into the mutagenesis vector pGJJ055 via BamHI/HindIII to create kinase-specific templates pDMS005 (Fyn), pDMS006 (Lck), and pDMS007 (Src). Comprehensive saturation mutagenesis was performed using the one-pot method with NNK primer sets designed for each kinase domain. Following electroporation into TOP10 electrocompetent cells and overnight growth, mutant libraries were harvested by plate scraping, and plasmid DNA was prepared by midiprep. BamHI/HindIII kinase fragments were cloned into pDMS002 to generate phospho-PCA libraries containing the Fyn substrate. Libraries were linearized at the barcode landing pad (AvrII/BsiWI) and barcoded using IDT oligonucleotide cassettes with NEBuilder Assembly. Each barcoded pool was bottlenecked to approximately 3% of transformants to maintain library diversity while ensuring adequate representation. To generate the complete 3×3 kinase-substrate panel, substrate cassettes were exchanged using AvrII/BamHI digestion and ligation with fragments from pDMS003 and pDMS004 to assemble barcode-[Fyn/Lck/Src]Mut-[fynsub/lcksub/srcsub], yielding nine phospho-PCA libraries (see Figure S12 for a construct map). Matched abundance-PCA libraries were constructed by ligating kinase-substrate-barcode cassettes into pGJJ045 via BsiWI/HindIII to generate barcode-[Fyn/Lck/Src]Mut-[fynsub/lcksub/srcsub]-noSH2.

#### Selection of barcoded biosensor libraries with methotrexate

BY4742 yeast cells were prepared for electroporation by growth to mid-log phase (OD_600_ ≈ 1.6), followed by washing with ice-cold water and electroporation buffer. Cells were incubated in 0.1 M LiAc/10 mM DTT for 20 minutes at 30°C before final preparation. Libraries (0.3 µg) were electroporated with salmon sperm DNA (50 µg) using standard parameters (2.5 kV, 25 µF, 200 Ω). Transformed cells were rescued in sorbitol/YPD medium and expanded in SC-URA medium for 48 hours. Following initial selection on uracil-deficient media, input samples were grown for approximately 4 generations in SC-URA/MET/ADE medium. Competition experiments were initiated by transferring cells into medium containing 200 µg/mL methotrexate at OD_600_ = 0.05 and growing for approximately 5 generations. For each library, 12-16 electroporation reactions were pooled across two biological replicates, requiring 1.35 L of medium per replicate. Cell pellets were harvested by centrifugation and stored at -20°C for subsequent plasmid extraction using ZymoPrep II columns.

#### Library preparation for next generation sequencing

Barcode regions were amplified using a two-step PCR protocol. In PCR1, eight parallel 50 µL reactions per library were performed using KAPA HiFi polymerase with barcode-flanking primers for 15 cycles at 65°C annealing temperature. Products were pooled and purified using DNA Clean & Concentrator-5 columns. PCR2 appended Illumina adapter sequences through four 50 µL Q5 HotStart reactions with index primers for 10 cycles. Final libraries (225 bp) were sequenced on Illumina platform with paired-end 50 bp reads targeting 50 million reads per library.

For barcode-variant mapping, the contiguous kinase-substrate-barcode segment (1.7 kb) was amplified using KAPA HiFi polymerase with 24 parallel reactions per kinase to preserve library complexity. Thermocycling parameters included 14 cycles with 10-second annealing at 71°C and extension at 20 seconds per kilobase. Products were pooled, purified, and sequenced on the PacBio Revio platform, generating approximately 45 million HiFi reads across four SMRT cells.

### Quantification and statistical analysis

#### Next generation sequencing analysis

We developed bioinformatics pipelines to construct barcode-variant look-up tables from long-read PacBio sequencing and measure barcode abundances in selection experiments from short-read Illumina sequencing.

##### Barcode-variant look up table construction

The goal of our analysis was to create a high-fidelity map between barcodes and their associated kinase variants from our long-read PacBio sequencing data. Due to sequencing errors, multiple reads from the same barcode may differ slightly. We sought to determine the true underlying sequence for each barcode by using Bayesian statistics to integrate information across all reads sharing that barcode. The first step in our analysis was to extract barcode and kinase variable region sequences from the PacBio reads. HiFi FASTQ reads from each kinase library were filtered for the exact vector context, the presence of both the DHFR–substrate cassette and a 3’ overhang motif. Reads with either anchor missing were discarded. Reads that passed this QC step were parsed to extract the landing-pad barcode and the kinase ORF. We validated the kinase library identity via a two-base tag embedded in the barcode (AG, CT, and TA for Fyn, Lck, and Src, respectively); only tag-consistent reads were retained. ORFs were length-checked (765 nt), translated, and compared codon-wise to the kinase-specific wild type to perform nucleotide and amino-acid level variant calling. All reads were grouped by their barcode identifier, creating sets of reads that originated from the same original molecule. Each barcode group contains multiple, noisy observations of the same “true” underlying sequence.

The next step in our analysis was to estimate the consensus variant sequence for each barcode using Bayesian statistics. For each position *i* in the variable region, we want to determine which nucleotide *b_i_* ∈ {*A, T, C, G*} is most likely the true base, given our observed data *D_i_*(e.g., all bases observed at position *i* in all reads with the same barcode). Bayes rule gives us

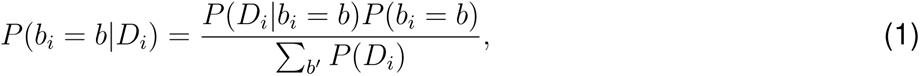

where *P* (*b_i_* = *b*|*D_i_*) is the posterior probability that the true base at position *i* is *b* given the observed data, *P* (*D_i_*|*b_i_* = *b*) is the likelihood of observing our data if the true base were *b*, which we take to be 0.9 for the correct base and 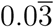 for errors. *P* (*b_i_* = *b*) is the prior probability that position *i* contains base *b* (which we set to 0.25 for a uniform prior), and *P* (*D_i_*) is a normalization constant (equal to Σ*_b_*_′_*P* (*D_i_*|*b_i_* = *b*^′^)*P* (*b_i_* = *b*^′^)) to ensure the probabilities sum to 1. To compute the position-wise consensus, at each position *i* we select the base with the highest posterior probability, e.g., 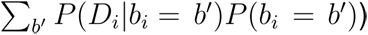. We use the maximum posterior probability at each position as a confidence score; an overall score for the complete consensus sequence was determined by computing the joint posterior probability assuming position-wise independence, *P* (consensus sequence correct) = 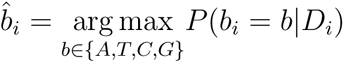. To ensure the accuracy of our lookup table, barcode-variant pairs with a joint posterior probability below 0.99 were filtered out and excluded from subsequent analyses.

##### Bioinformatic analysis of deep mutational scanning data

Our bioinformatics pipeline used MAGeCK (Model-based Analysis of Genome-wide CRISPR-Cas9 Knockout) to compute per-barcode log_2_ fold changes for outlier detection, then applied Rosace’s Bayesian hierarchical model to the filtered raw count matrix to generate fitness scores that account for barcode-to-barcode variability. Paired-end short-read sequencing data from selection experiments were processed using the MAGeCK pipeline to count barcodes. Paired-end FASTQ files were quality-filtered and merged using fastp, with a minimum overlap of 6 base pairs. Barcode abundance was quantified using MAGeCK count, with 34 nucleotides trimmed from the 5’ end to remove constant upstream sequences. This analysis generated sample-labeled count matrices for both pre-selection and post-methotrexate selection replicates. For each screening condition, enrichment was assessed with MAGeCK ‘test‘ by comparing pre- and post-selection replicates, using the –remove-zero flag. This yielded per-barcode log_2_ fold changes and false discovery rate adjusted p-values at the single-barcode level. To link Illumina barcode counts with PacBio-derived barcode-variant assignments, MAGeCK outputs were processed through several quality control steps. True barcodes were extracted as sequences 5’ of the constant flanking anchor (GCCTAGGCAGCTATGACCATGATTACGCCAAGCG) and filtered by expected library tags (AG, CT, or TA for Fyn, Lck, and Src libraries, respectively). Only barcodes with minimum pre-selection coverage (>20 reads) were retained for analysis. Barcode sequences were matched to the PacBio lookup table using exact sequence identity. Analysis was restricted to barcodes corresponding to two or fewer amino acid changes. To remove outlier measurements, barcode-level log_2_fold changes were z-score normalized within each variant group, and barcodes with |z| > 2 were excluded from further analysis. Final variant matrices were generated by collapsing barcode counts across wild-type, single-nucleotide synonymous, and single-amino-acid variants. These matrices were then analyzed with Rosace to generate fitness scores for each variant. The resulting 18 screening libraries achieved a barcode-variant coverage of 10-25X and included between 95.9% and 99.3% of all possible single amino acid mutations within the 254-residue kinase domain (Table S1).

#### Abundance-PCA score normalization and enrichment analysis

Raw abundance scores for each kinase-mutation pair were normalized to account for systematic variation across kinases and substrates. To enable cross-kinase comparison, scores were scaled using a relative nonsense baseline: the mean abundance score of all nonsense (stop-gain) variants was computed for each kinase, and a scale factor was applied to align these nonsense means to a common reference value. Synonymous variant scores were used to zero-center the distribution, anchoring functionally neutral changes at zero. For each kinase, variant-level abundance scores were averaged across the three substrate backgrounds to yield a single score per mutation. These normalized abundance scores were used to identify residue positions enriched for loss-of-function (LoF) or gain-of-function (GoF) variants. Variants were preliminarily classified as GoF if their normalized abundance score exceeded the mean of all positive scores for that kinase, and as LoF if the score fell below 0.45-fold the mean of all negative scores. These thresholds were selected to reflect consistent phenotypic patterns across kinases. For each residue position, a one-sided Fisher’s exact test assessed whether LoF or GoF variants were significantly over-represented at that position relative to all other positions. Benjamini-Hochberg correction was applied across all tested positions, and residues with an adjusted false discovery rate (FDR) ≤ 0.10 were designated as LoF- or GoF-enriched positions for the abundance phenotype.

#### Solvent-Accessible Surface Area (SASA) computation and stratification

To assess the relationship between surface exposure and mutational tolerance, we computed per-residue solvent-accessible surface area for the catalytic kinase domains of three Src-family kinases using crystal structures: Fyn (PDB: 2DQ7), Lck (PDB: 3LCK), and Src (PDB: 1YI6). Only polymeric protein atoms were considered in the analysis. Per-atom SASA was computed in PyMOL using a routine modeled on the Lee-Richards/Connolly surface algorithm. We isolated polymeric protein atoms, enabled solvent dot repre-sentation (dot_solvent=1 with a 1.4Åprobe radius), and computed per-atom SASA using cmd.get_area(…, load_b=1) to store accessibility values in the B-factor field. Atoms with SASA < 2.5 Å^2^ were removed, and the remaining atoms were saved as exposed. Residues were classified as surface-exposed if any constituent atom exceeded the 2.5 Å^2^ cutoff via byres expansion; otherwise, they were labeled non-exposed. To enable stratified analysis of mutational effects, we assigned ordinal exposure categories to each aligned position based on per-residue SASA: strongly exposed (SASA > 150 Å^2^, label = -1), moderately exposed (50 Å^2^ < SASA ≤ 150 Å^2^, label = 0), buried (10 Å^2^ ≤ SASA < 50 Å^2^, label = 1), strongly buried (2.5 Å^2^ ≤ SASA < 10 Å^2^, label = 2), and core/hydrophobic (SASA < 2.5 Å^2^, label = 3). This stratification allowed us to test whether the structural context of a residue—particularly its degree of solvent exposure—predicts the functional impact of mutations as measured by abundance-PCA.

#### Decoupled activity score harmonization and enrichment analysis

Variant-specific activity effects decoupled from abundance were extracted from the hierarchical Bayesian model for each kinase-position-mutation combination. To enable cross-kinase comparison while preserving the interpretability of zero as a neutral baseline, we applied a two-step normalization procedure. First, within each kinase, positive and negative scores were rescaled independently: positive values were divided by the kinase-specific mean of all positive scores, negative values were divided by the absolute value of the kinase-specific mean of all negative scores, and zero values remained unchanged. Second, a between-kinase offset correction was applied by aligning the mean decoupled score of high-confidence positive variants (score > 0.20) to a common reference, ensuring kinases shared a comparable scale for gain-of-function effects.

Normalized decoupled activity scores were converted to categorical variant calls using fixed thresholds: GoF if score > 0.24, LoF if score < -0.21, and neutral otherwise. These thresholds were applied uniformly across all kinases. To identify positions enriched for LoF or GoF variants, we performed a one-sided Fisher’s exact test for each residue position (1–254) in each kinase, comparing the proportion of LoF (or GoF) variants at that position to all other positions. P-values were corrected for multiple testing using the Benjamini-Hochberg procedure across all residue positions, and positions with FDR ≤ 0.10 were designated as significantly enriched.

To define conserved regulatory positions across kinases, we required that a residue be significantly enriched in at least two of the three kinases. To recover positions where two kinases showed significant enrichment but the third narrowly failed the FDR threshold, we applied a complementary high-stringency criterion: if the third kinase had ≥ 2 variants at that position falling in the bottom 20th percentile of decoupled activity scores (for LoF) or the top 20th percentile (for GoF), the position was designated as enriched for that kinase as well, yielding a consistent cross-kinase annotation.

#### Sampling and analysis of kinase conformational ensemble with BioEmu

To understand how gain-of-function mutations alter kinase activity at a structural level, we used BioEmu to sample conformational ensembles for wild-type and mutant kinase domains. BioEmu is a recently developed generative deep learning method that enables scalable emulation of protein equilibrium ensembles, providing a computationally efficient alternative to traditional molecular dynamics for exploring conformational landscapes. For each variant of interest, we generated a mutant sequence by substituting the target amino acid at the specified position and used BioEmu to sample 4,000 conformations per sequence. Each sampled ensemble was output as a topology PDB file and a trajectory file, capturing the distribution of accessible conformational states. To ensure structural quality and accurate side-chain geometry, we reconstructed and relaxed side chains for all sampled models using BioEmu’s hpacker-openmm workflow. To classify conformational states across the relaxed ensembles, we used Kincore, a validated tool for annotating kinase activation states based on structural features. We iterated frame-by-frame through each trajectory using MDAnalysis, wrote temporary PDB snapshots, and invoked Kincore’s classifier to extract per-frame state labels (active DFG-in, inactive DFG-in, inactive DFG-out, or unfolded). This approach allowed us to quantify how mutations shift the equilibrium distribution of kinase conformational states, providing mechanistic insight into the structural basis of activity-biasing mutations identified through our deep mutational scanning experiments.

#### Interaction parsing of kinase structures

To annotate specific non-covalent interactions present in BioEmu-sampled models, we analyzed each PDB snapshot in a standardized pipeline: Each PDB was protonated and assigned atomic charges via PROPKA 3 (for residue *pK_a_* estimation) and PDB2PQR (Amber force-field parameters), producing PQR files for consistent electrostatics. These PQR files were used as inputs to PROPKA; the generated Coulomb table was then parsed to list salt-bridge partners of each captured residue, treating them as putative salt-bridge/electrostatic pairs. Side-chain and backbone H-bond partners reported by PROPKA were captured separately.

#### Evolutionary coupling analysis of position 195

To identify the network of co-evolving residues involving residue 195, we first computed long-range coevolution scores for Fyn, Lck, and c-Src. CSV outputs of long-range coupling scores were loaded per kinase and merged into a single table. We constructed an undirected graph with residue indices as nodes and EV-coupling probabilities as edge weights (NetworkX). From this graph, we extracted the local coevolutionary neighborhood of position 195 by thresholding edges at high confidence (weight > 0.95) and identifying a co-evolving cluster conserved across three kinases.

#### Protein language model analysis of allosteric sites

To evaluate whether protein language models could identify functionally important allosteric sites in kinase domains, we computed ESM-1b representations and attention patterns for the kinase-domain sequences of Fyn, Lck, and c-Src using the FAIR-ESM Torch Hub model esm1b_t33_650M_UR50S in evaluation mode. For each sequence, we performed a forward pass requesting all 33 layers (repr_layers=1-33) with return_contacts=True, extracted per-residue representations from the final layer, and aggregated the model’s attention heads into a mean residue × residue attention matrix after removing special tokens (BOS/EOS). We defined an a priori active-site set covering known catalytic and recognition motifs, including *β*3-Lys, *α*C-Glu, the HRD/DFG/APE motifs, P-loop residues, and the activation-loop tyrosine with flanking residues. For each kinase, we derived two per-residue attention vectors: (i) global attention, computed as the column-wise mean of the full attention matrix, and (ii) active-site attention, computed as the column-wise mean after subsetting rows to active-site residue indices. Putative allosteric residues were identified using an active-site attention threshold of 0.005 with an adjacency exclusion that removed active-site residues and their ±1 sequence neighbors to focus on distal coupling. Mean attention matrices were serialized for downstream analysis, and visualizations overlaid global and active-site attention traces across residue positions with vertical markers indicating active-site locations and a dashed horizontal line marking the 0.005 threshold.

#### Molecular dynamics simulations

Kinase-substrate complex structures were generated using Chai-1 in multimer mode with ligands and geometric constraints enabled, where each complex contained one kinase domain (Chain A), one substrate peptide (Chain B), two magnesium ions (Chains C and D), and ATP (Chain E), with MMSeqs2 and template matching settings applied. To ensure catalytically relevant conformations, we implemented specific geometric constraints including inter-chain contacts (4.0 Ådistance) between A:R119 and B:Y8, and between A:D120 and B:Y8 to position the phosphorylatable tyrosine near the HRD motif, as well as pocket constraints (4.0 Ådistance) between ATP (Chain E) and B:Y8, and between ATP and A:L127 to maintain proper positioning within the catalytic cleft. For molecular dynamics simulations, the coordinates of ATP and Mg^2+^ were refined by superimposing reference crystallographic structures onto the Chai-1 predicted models using PyMOL. The ATP positioning was based on the nucleotide pose in PDB structure 1QMZ, and the Mg^2+^ coordinates, which were absent from 1QMZ, were derived from PDB structure 1IR3. Molecular dynamics simulations were performed using GROMACS 2025.2 with the CHARMM36-jul2022 force field and the TIP3P water model. Protein topologies and position restraints were generated using pdb2gmx. The simulation protocols included energy minimization, NVT equilibration, NPT equilibration, and 500 ns production runs under standard conditions. Trajectories were processed using trjconv to center molecules and maintain molecular integrity (-pbc mol -center -ur compact) and to remove periodic boundary condition artifacts (-pbc nojump), with additional PBC correction for individual protein chains performed using the MDAnalysis Python package when GROMACS processing was insufficient, and stable trajectory segments of 100 ns duration were identified based on root mean square fluctuation (RMSF) analysis and visually inspected using ChimeraX for selection as representative conformational states.

#### Molecular dynamics trajectory analysis

To analyze the structural trajectories generated by our molecular dynamics simulations, we analyzed the structural fragment correlations and residue interaction graphs. The details of both analyses are described below.

##### Structural alphabet encoding with AllohubPy

To identify fragment pairs with differential interactions between wild-type and mutant kinases, MD trajectories were converted to structural alphabet representations using AllohubPy’s encoder (alphabet: M32K25, block size = 100, with block randomization). Each system was analyzed as a two-condition comparison with two replicates per condition. For each fragment pair, we computed log_2_ fold changes (mutant vs. wild-type) and adjusted p-values (Benjamini-Hochberg FDR). Significant fragments were identified using FDR <10^−4^ and variant-specific fold-change thresholds (up-regulated: log_2_FC >4; down-regulated: log_2_FC <-4). Fragment definitions were extended by one residue downstream for graph construction.

##### Residue interaction graph construction and pathway analysis

For each replicate, we calculated per-frame interaction frequencies at both residue and atom levels. Mutant and wild-type replicate means and their differences were computed for all residue pairs. To enforce replicate concordance, we applied a minimal frequency filter that required either the wild-type or mutant mean frequency to be greater than 0.01. Differential interactions were identified using variant-specific mean difference thresholds (P203R: 0.115; Q65D: 0.3) that were determined by balancing two objectives: (1) capturing local contact changes in the immediate vicinity of the mutation site, and (2) maintaining global stringency to identify the most substantial allosteric perturbations. The threshold for each variant was set as the minimum value that retained clear differential contacts within the first shell of residues surrounding the mutation site while filtering noise from more distal regions. P203R required a lower threshold (0.115) because this mutation produced more subtle, distributed contact changes compared to the larger local perturbations induced by Q65D (0.3). To capture conserved communication pathways, we identified high-frequency, stable contacts (average wild-type and mutant frequencies both >0.8 with a difference <0.1) and included neighboring residues as potential relay points.

Residue contact networks were constructed with residues as nodes and frequent contacts as edges, colored by interaction changes (red: up-regulated in mutant; blue: down-regulated; gray: conserved; dashed gray: neighboring connections). Only connected components with ≥ 4 nodes were retained. For P203R, we defined important nodes as the mutation site, all substrate residues with direct kinase contacts, and residues within differential AllohubPy fragments. We then reduced the full graph *G* to a minimal subgraph *H* that: (i) preserves all important residues as individual nodes, (ii) collapses intervening intermediate residues with conserved interactions into supernodes, and (iii) retains only the minimum set of intermediates required to maintain connectivity between important residues, thereby identifying the shortest allosteric communication pathways. For E113H, we compared interactions across multiple metastates by binarizing contact frequencies (threshold: 0.3) and computing intersections across selected metastate sets.

##### Identification of specificity-determining variants

During the development of our hierarchical Bayesian model, we observed that the specificity score was approximately an order of magnitude smaller than the other scores. This lower score reduced the model’s ability to confidently distinguish true directional effects, resulting in increased uncertainty in the local false-sign rate^110^. To ensure consistent statistical rigor across all mutation effect scores in our multi-dimensional inference framework, we applied a uniform significance threshold strategy. For specificity scores, we required that mutations satisfy both a 94% credible interval test (ensuring the posterior probability of a directional effect is ≥97%) and a Kolmogorov-Smirnov test comparing the posterior distribution to the prior (ensuring meaningful divergence from the prior distribution)^111,112^. This dual-threshold approach guards against cases where the credible interval criterion might be satisfied due to a narrow posterior that has shifted only modestly from the prior, without substantial evidence accumulation. The KS test ensures that the posterior distribution has genuinely moved away from the prior, providing an additional layer of confidence that the observed specificity effect reflects true information content in the data rather than minor distributional shifts^112^.

## Additional resources

Code for analysis pipelines and processed datasets are available at the URLs specified in the Data and Materials Availability statement. Plasmids are available through Addgene.

## Supplementary materials

Supplementary Text

Figs. S1 to S13

Tables S1 to S3

Movie S1

Data S1

