## Supplemental Information for "Uncovering the molecular basis of kinase activity and substrate recognition with phospho-PCA"

**This PDF file includes:**

Supplementary Text
Figures S1 to S13
Tables S1 to S3
Captions for Movies S1
Captions for Data S1-S3

**Other Supplementary Materials for this manuscript:**

Movies S1
Data S1-S3

### Supplementary Text

#### Decomposition of phospho-PCA data through hierarchical Bayesian modeling

Deep mutational scanning (DMS) experiments often confound several biological effects that - in an ideal world - should be carefully separated to extract meaningful insights. In our kinase studies, measured phosphorylation depends on protein folding stability, the probability the kinase is in an active state, and the specificity of the kinase-substrate interaction. Since protein misfolding reduces kinase abundance, which in turn decreases phosphorylation, folding stability acts as a confounder that must be separated from the other two effects. This is necessary to distinguish between mutations that genuinely alter substrate recognition or kinase activity versus those that simply impair overall protein stability.

Our experimental design uses a dual DMS screen to address this confounding issue: abundance-PCA measures kinase folding and stability (i.e., the confounding factor), while phospho-PCA measures the kinase-substrate interaction (i.e., the target signal plus the confounder). Our experimental data consists of abundance-PCA and phospho-PCA scores for each kinase identity ($i$), substrate identity ($j$), position along the kinase domain ($k$), and amino acid substitution ($l$). To separate these effects, we developed a hierarchical Bayesian regression model that decomposes the observed phospho-PCA signal into three distinct components:

$$\text{phospho-PCA}_{ijkl} = a_{ij} \times \text{Abundance}_{ikl} + b_{ij} + \text{Activity}_{ikl} + \text{Substrate specificity}_{ijkl}, \qquad \text{(S1)}$$

where each term captures distinct biological effects operating at different hierarchical levels of organization. We note that the abundance-PCA scores are averaged over all 3 substrates, yielding $\text{Abundance}_{ikl}$. We further note that we add the constraint $\sum_j \text{Substrate specificity}_{ijkl} = 0$ to ensure identifiability. This model architecture reflects the nested structure of biological variation across three levels, each addressing distinct sources of experimental and biological variation. These levels are given below.

- **Level 1: Kinase-substrate experiment effects.** At this level, we group mutations by each unique kinase-substrate pair ($i$ and $j$) to capture overall biochemical context and experimental batch effects. This level also posits a linear relationship between abundance- and phospho-PCA scores. There are two key parameters at this level - $a_{ij}$, a slope parameter capturing how abundance affects phospho-PCA scores for each specific kinase-substrate pair, and $b_{ij}$, an intercept representing the baseline phosphorylation activity independent of abundance effects.

- **Level 2: Kinase activity effects.** The second level addresses kinase activity effects by grouping mutations according to kinase identity, mutation position, and amino acid change ($i$, $k$, and $l$). This captures changes in kinase activity caused by mutations that are independent of substrate identity. The corresponding model term represents mutation-induced perturbations to kinase catalytic activity that affect all substrates equally, reflecting changes to the fundamental enzymatic machinery rather than substrate-specific recognition elements. For example, this term captures mutations that introduce a bias towards active or inactive conformations.

- **Level 3: Kinase-substrate specificity effects.** At the finest resolution, we group by all four factors - kinase ($i$), substrate ($j$), position ($k$), and mutation ($l$) to capture effects where mutations differentially impact particular kinase-substrate pairs. This accounts for potential substrate-specific modulation of a mutation's impact, and represents the true substrate recognition signal we aim to isolate from confounding abundance effects.

Our model design was guided by our intuition regarding which aspects of the experiment should affect which model terms. The abundance score quantifies how mutations alter the abundance of the kinase itself; since phosphorylation activity is inherently influenced by kinase concentration, failing to account for these changes would confound our measurement of kinase activity. Our model addresses this by incorporating abundance as a covariate with kinase-substrate specific slopes ($a_{ij}$), allowing each kinase-substrate pair to have its own abundance-phosphorylation relationship. This should (in principle) statistically separate true changes in enzymatic activity from abundance-driven changes, enabling more precise inference of the functional impact of mutations. We empirically observed little variance in the abundance-PCA scores for each kinase across the three substrates. The activity score was assumed to be conserved across multiple substrates for a single kinase, adding additional regularization to further enhance the robustness of our inference.

To regularize parameter estimates while allowing the data to drive model conclusions, we employ weakly informative priors. Specifically, we define:

- **Abundance effect slopes** $a_{ij} \mid \sigma_a \sim \mathcal{N}(0, \sigma_a^2), \sigma_a \sim \mathrm{HalfNormal}(1)$.
- **Baseline activity intercepts.** $b_{ij} \mid \sigma_b \sim \mathcal{N}(0, \sigma_b^2), \sigma_b \sim \mathrm{HalfNormal}(1)$.
- **Kinase activity effects.** $f_{ikl}^{(\text{activity})} \mid \sigma_{\text{activity}} \sim \mathcal{N}(0, \sigma_{\text{activity}}^2), \sigma_{\text{activity}} \sim \mathrm{LogNormal}(\log 0.5, 0.5^2)$.
- **Substrate specificity effects (with constraint).** We define

  $$C_{i,k,l} = \{(i, j', k, l) \text{ present in the data for fixed } (i, k, l)\}.$$

  For each activity parent $(i, k, l)$, the set of its specificity children is

  $$\delta_{ijkl}^{(\text{specificity})} \mid \sigma_{\text{specificity}} \sim \mathcal{N}(0, \sigma_{\text{specificity}}^2), \sigma_{\text{specificity}} \sim \mathrm{LogNormal}(\log 0.25, 0.5^2).$$

  Again, we additionally constrain $\sum_{(i,j',k,l)\in C_{i,k,l}} \delta_{ij'kl}^{(\text{specificity})} = 0$.

Observation level sampling, where $f_{ikl}^{\text{abundance}}$ is the abundance-PCA data for $(i, k, l)$ is given by the following equations:

$$\begin{aligned} \mu_{ijkl} &= a_{ij} f_{ikl}^{\text{abundance}} + b_{ij} + f_{ikl}^{(\text{activity})} + \delta_{ijkl}^{(\text{specificity})}, \\ y_{ijkl} \mid \mu_{ijkl}, \sigma &\sim \mathcal{N}(\mu_{ijkl}, \sigma^2), \text{with } \sigma \sim \mathrm{LogNormal}(\log 0.5, 0.5^2) \end{aligned}$$

We also note the following additional model details.

**Partial pooling and non-centered parameterization.** We use z-score normalization for all group effects - specifically, we use $X = \sigma Z$ with $Z \sim \mathcal{N}(0, 1)$ for group effects $a_{ij}$, $b_{ij}$, $f_{ikl}^{(\text{activity})}$, and $\delta_{ijkl}^{(\text{specificity})}$. We found that this yielded shrinkage toward a common mean (partial pooling), improved posterior geometry by separating scale from standardized randomness, helped mitigate Neal's funnel, and reduced divergences.

**Nested-deviation + sum-to-zero removes the ridge.** Specificity effects were modeled as mean-zero deviations within each activity parent: $\sum_{(i,j',k,l)\in C_{i,k,l}} \delta_{ij'kl}^{(\text{specificity})} = 0$. This makes $f_{ikl}^{(\text{activity})}$ the identifiable parent mean and the $\delta_{ijkl}^{(\text{specificity})}$ identifiable contrasts, eliminating the additive trade-off between parent and children.

The model was implemented in PyMC and fitted to our data using the No-U-Turn Sampler (NUTS). Convergence statistics (e.g., $\hat{r}$ and effective sample size) were used to ensure sample quality.

### Phospho-PCA reveals the limitations of protein language models

Recent work suggests that protein language models (pLMs) can identify functionally important allosteric sites by identifying distal sites that exhibit high attention scores with known active sites[113,114]. The data generated in our study, in particular the decomposed activity scores, provide an opportunity to test how well this approach works for protein kinases. Following these recent studies, we used the protein language model, ESM-1b, to compute attention scores between each residue and a set of active site residues for Fyn, Lck, and c-Src and compared sites with high scores - putative allosteric sites - with our kinase activity scores. The results are shown in Figure S3; a detailed description of our analysis is given in the Methods section. We found that ESM-1b was able to identify 10 candidate high-attention sites across these three kinases; eight of these corresponded to variants significant differences in kinase activity scores in our data. These positions overlapped with loss-of-function variants we identified at positions 53, 71, 73, 112, and 180 as well as the conserved gain-of-function variant at position 236. However, we noted that ESM-1b identified two sites - 125 and 162 - that were adjacent to positions 126 and 161 that displayed significant activity effects in our data, suggesting that ESM-1b may sometimes capture variants that are only in the general vicinity of functionally important regions. In addition, ESM-1b's analysis was incomplete, as 30 positions containing variants with significant activity scores but minimal impact on abundance were not flagged by ESM-1b. Compared to our data, ESM-1b attention had a recall of $\approx$0.21 and a precision of 0.8, indicating that pLMs miss a substantial fraction of functionally important allosteric sites. This may also reflect a fundamental limitation of self-supervised learning on sequence data alone, as co-evolutionary information is distinct from the allosteric communication networks that emerge from protein dynamics.

### Specificity determining variants

In this section, we describe the collection of variants identified by our previously described bioinformatics pipeline, which impact substrate specificity.

- *Fyn*: Our analysis revealed several specificity-determining mutations in Fyn. The S123L mutation decreased activity toward the Fyn substrate by replacing a small polar serine with a hydrophobic leucine, demonstrating that precise physicochemical properties at surface-exposed positions contribute to substrate recognition. Additionally, position 248 is conserved between Fyn and Lck but differs for c-Src, further supporting its role in substrate specificity determination.

- *Lck*: Lck exhibited the largest collection of specificity-determining mutants, which may reflect its distinct evolutionary position among SFKs[115]. The A9Q and A9S mutations decreased specificity toward the Lck substrate while increasing specificity toward the Fyn substrate by exchanging a nonpolar amino acid for a polar one. The F19T mutation consistently decreased specificity toward the Lck substrate, indicating that the native aromatic residue is essential for optimal substrate recognition. The Q33P mutation reduced the phosphorylation of the Lck substrate by replacing a flexible polar residue with proline, thus introducing structural constraints that likely impair the conformational flexibility required for substrate binding. Position 43 shows extensive specificity switching: in Lck, the A43S and A43T mutations reduced specificity toward the Lck substrate. These mutations, adjacent to the critical E44 substrate-binding residue, introduce a polar hydroxyl group capable of hydrogen bonding. The Q65D mutation has dual effects on local electrostatics: it increases specificity toward the Fyn substrate by introducing a favorable negative charge while reducing activity toward the Lck substrate, likely due to disruption of Lck-specific electrostatic interactions. The I90H mutation reduced activity toward the Lck substrate by substituting a buried hydrophobic isoleucine with a polar histidine. The E113K and E113H variants, which exchange a negative charge for a positive charge, showed reduced specificity toward the Lck substrate and increased scores towards Fyn and c-Src substrates. The adjacent R114Y mutation displayed a similar pattern, with decreased specificity towards the Lck substrate by replacing a positively charged arginine with a bulky aromatic tyrosine. The opposite charges present at position 113 between Lck and c-Src/Fyn, and the importance of this location for the preference of Lck substrate, suggest that this residue could be one of the important divergent residues separating the SrcA and SrcB subfamilies of SFKs. The T172R and T172K mutations enhanced activity toward the Fyn substrate while reducing Lck-specific activity by introducing a positively charged side chain that may reinforce electrostatic complementarity. We also observed that the P203H and P203R mutations, which introduce a positive charge, reduced activity toward the Lck substrate. This convergence implies that position 203 acts as a key gating residue controlling substrate discrimination between Lck-like and Fyn/Src-like preferences. This observation was consistent with previous studies on SFKs[22]. We further explored the impact of this location in our molecular dynamics study. Last, the R247Q mutation reduced activity toward the Lck substrate by neutralizing a positive charge.

- *c-Src*: c-Src contained a collection of mutations that highlight the importance of surface charge and polarity. The mutation T19F drove preferential phosphorylation of the Lck substrate, complementing the Lck F19T findings and confirming that aromatic versus polar residues at this position provide a specificity switch between these kinases. At position 43, the Q43E mutation increased activity toward the Fyn substrate by introducing a negative charge that alters local electrostatics, while the Q43V mutation decreased Fyn activity but enhanced Lck activity by replacing a polar residue with a hydrophobic one. All three kinases have different amino acids with different properties at this site adjacent to E44, a critical residue for substrate binding and phosphorylation. The H226Y and H226W mutations increased activity toward the Lck substrate while decreasing activity toward the c-Src substrate by enhancing hydrophobic contact with the substrate, which complements the Leu +1 in the Lck peptide but clashes with the Asp +1 in the c-Src peptide. Position 248 revealed a polarity-dependent specificity switch, as both A248T and A248N mutations increased specificity toward the Fyn substrate while reducing specificity toward the c-Src substrate by introducing a polar residue at a previously hydrophobic site. Notably, position 247 is conserved between c-Src and Fyn, whereas position 248 is conserved between Fyn and Lck, suggesting these adjacent positions evolved complementary roles in substrate discrimination.

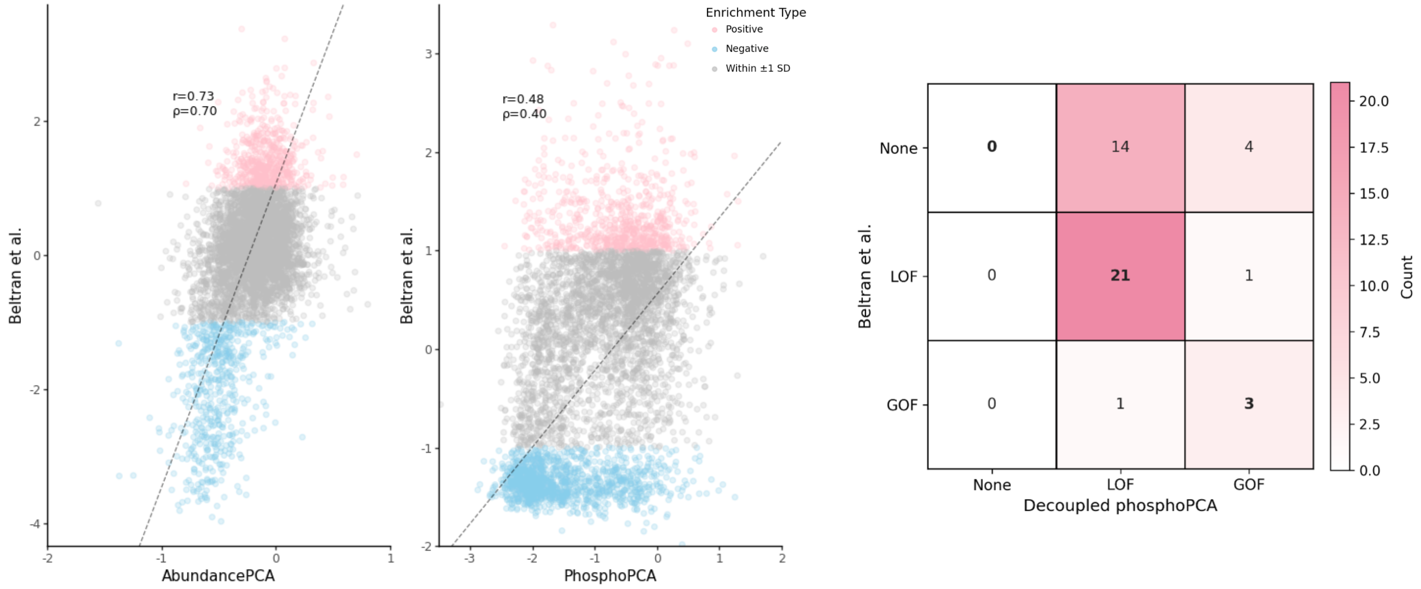


Figure S1: Validation of phospho-PCA measurements against published deep mutational scanning data. Left and middle panels: Scatter plots comparing abundance-PCA scores (left) and phospho-PCA scores (middle) from this study with corresponding measurements from Beltran et al. for c-Src kinase. Points are colored by enrichment type: positive (red), negative (blue), or within ±1 standard deviation (gray). Right panel: Contingency table showing overlap between loss-of-function (LOF) and gain-of-function (GOF) variants identified in decomposed kinase activity scores from phospho-PCA, compared to the categories from Beltran et al., demonstrating strong concordance between the two studies.

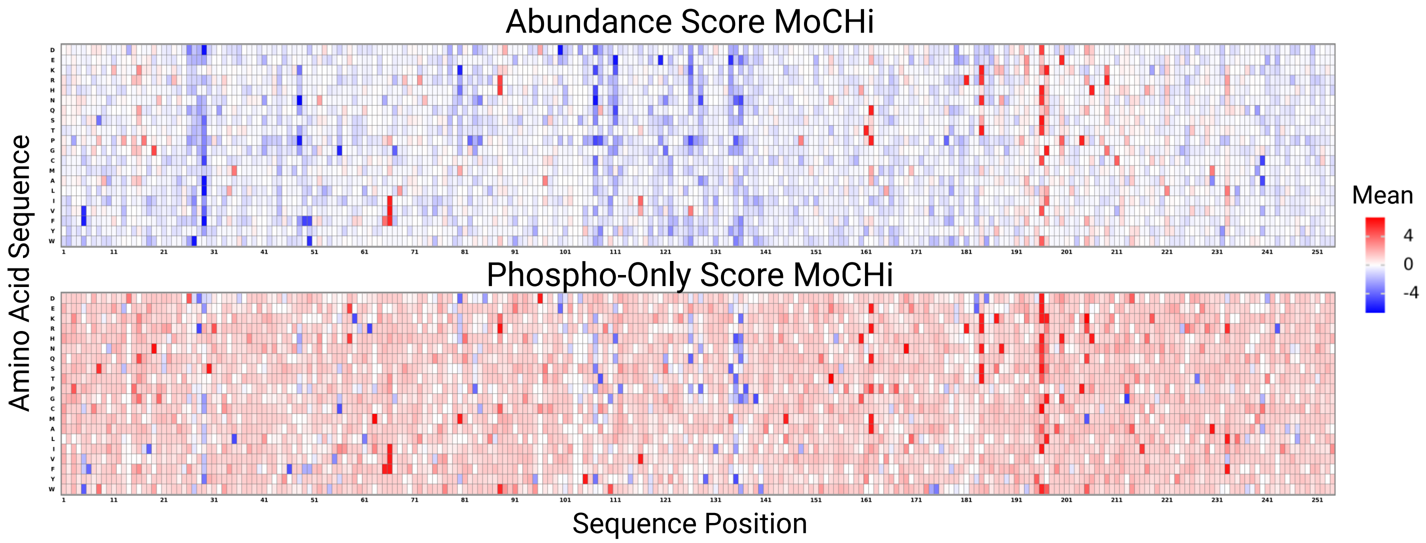


Figure S2: MoCHI model reveals incomplete decomposition of Fyn mutational data. Top panel: Abundance scores from MoCHI decomposition showing position-specific effects across the Fyn kinase domain. Bottom panel: Phospho-only (activity) scores from MoCHI showing residual mixed effects that are not fully separated from abundance contributions. The incomplete separation of abundance and activity effects motivated the development of our hierarchical Bayesian model, which better accounts for the nested structure of kinase mutational effects.

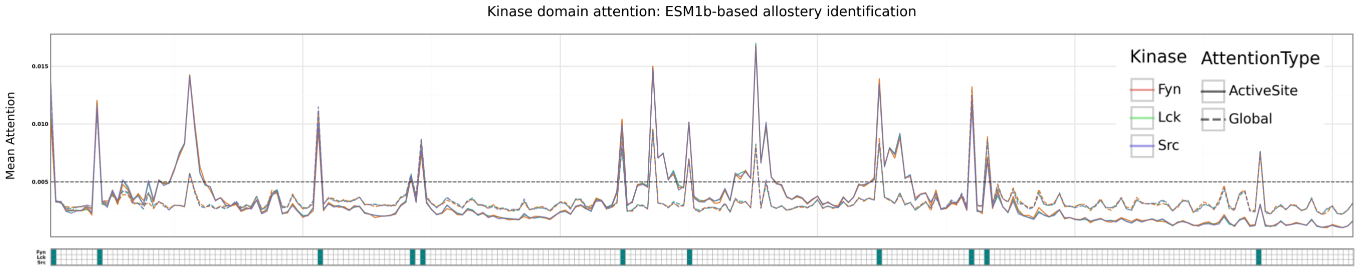


Figure S3: Protein language model attention analysis identifies a subset of functionally important sites. Top: Line plot showing ESM-1b attention scores across kinase domain positions for Fyn, Lck, and c-Src. Two attention metrics are shown: global attention (thin lines) and active-site-specific attention (bold lines). The dashed horizontal line indicates the threshold (0.005) for identifying putative allosteric sites. Vertical lines mark known active site positions. Bottom: Heatmaps showing the distribution of decomposed activity scores and amino acid properties at each position, allowing comparison between ESM-1b predictions and experimentally determined functional effects. High-attention sites show partial overlap with experimentally identified activity-biasing mutations. To generate this plot, we applied the allosteric site prediction pipeline from Kannan et al. (bioRxiv 2024, doi: 10.1101/2024.10.03.616547) to identify ten allosteric positions: 10 (G-loop, ATP binding), 53 ($\beta$4), 71 and 73 ($\beta$5, near gatekeeper residue), 112 (HE), 125 (catalytic loop), 162 (P+1 loop, substrate positioning), 180 and 183 (HF), and 236 (HH). Decoupled phospho-only scores show strong correspondence with ESM1b predictions, capturing conserved loss-of-function at positions 53, 71, 73, 112, and 180, and conserved gain-of-function at position 236. Notably, positions 126 and 161, immediately adjacent to predicted allosteric sites 125 and 162, also display decoupled activity effects. Most identified positions exhibit conserved phosphorylation activity effects, without corresponding changes in abundance.

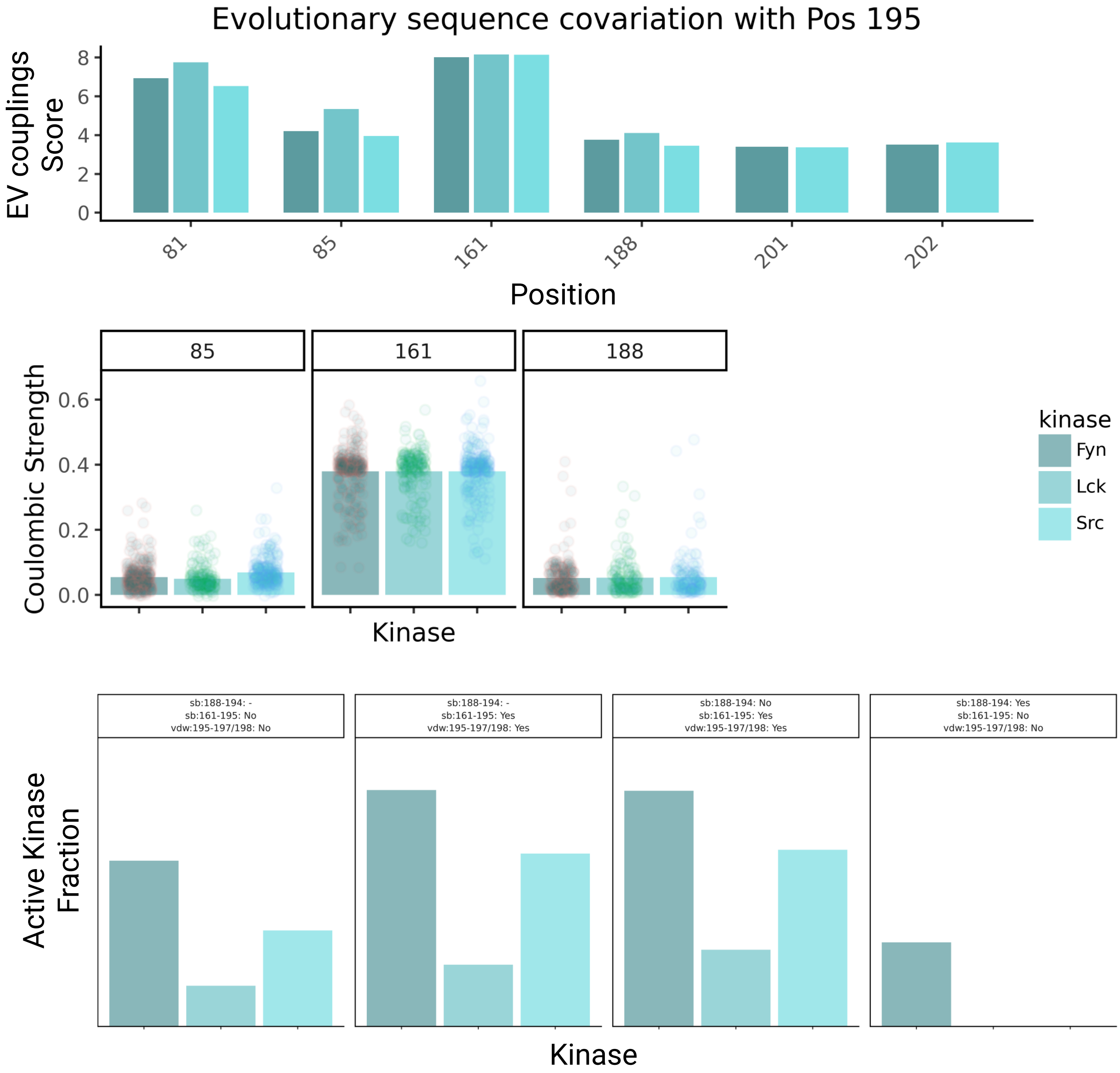


Figure S4: Evolutionary and biophysical characterization of the V195E gain-of-function mutation. Top: Bar plot showing evolutionary coupling scores between position 195 and other positions in the kinase domain, revealing a co-evolutionary network. Middle: Box plots comparing Coulombic interaction strengths between residue 195 and network partners (85, 161, 188) across wild-type and V195E mutant ensembles for Fyn, Lck, and c-Src. Bottom: Bar plots showing the fraction of active versus inactive conformations (classified by Kincore) for wild-type and V195E variants, stratified by presence or absence of the 161-195 salt bridge. The V195E mutation increases active conformation fractions when the 161-195 salt bridge is present.

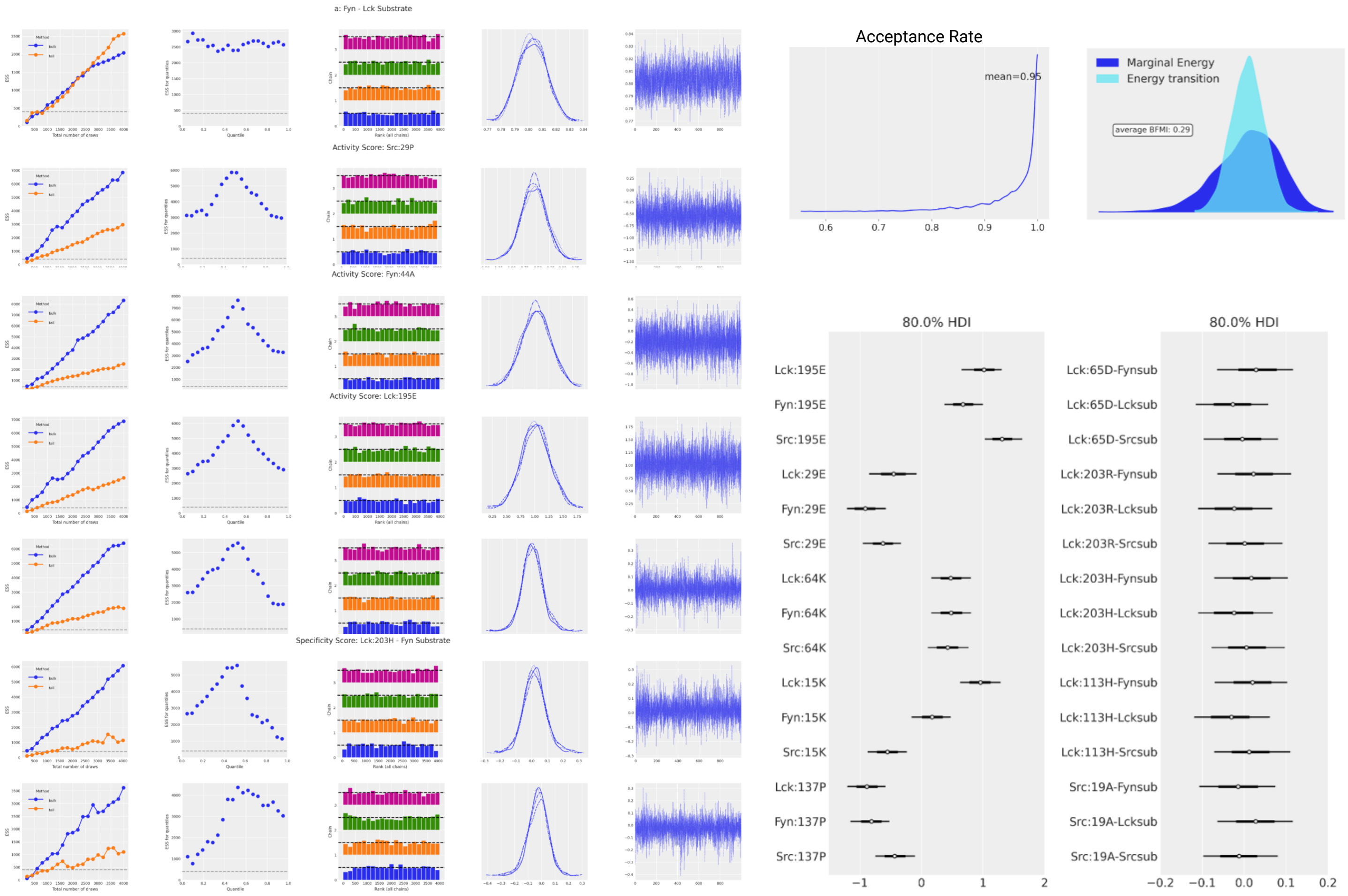


Figure S5: Hierarchical Bayesian model diagnostics demonstrate robust parameter inference. Left columns: Diagnostic plots for MCMC sampling including trace plots, rank plots, and energy plots for representative parameters at each hierarchical level (abundance scale factors, baseline activity, kinase activity effects, substrate specificity effects). Middle: Posterior distributions for selected parameters showing well-defined peaks and adequate sampling. Right top: Acceptance rate versus mean accept stat showing sampling efficiency. Right middle: 80% highest density intervals (HDI) for slope parameters across all kinase-substrate pairs. Right bottom: 80% HDI for selected activity and specificity parameters. All diagnostics indicate successful convergence with $\hat{r}$ values near 1.0 and adequate effective sample sizes.

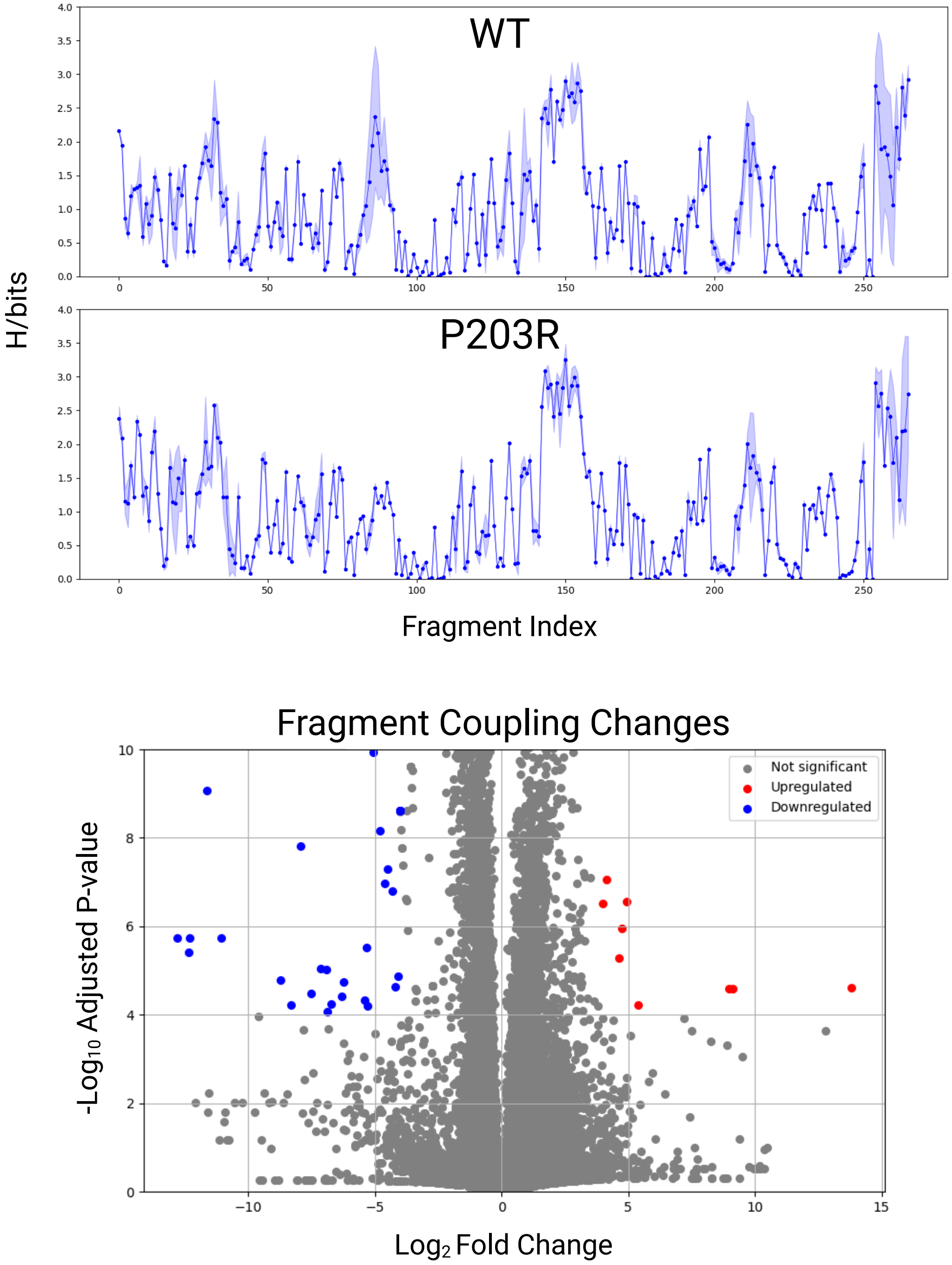


Figure S6: Allosteric hub analysis identifies differentially correlated protein segments in P203R mutant. Top: Positional entropy profiles comparing wild-type Lck and P203R mutant across molecular dynamics trajectories, showing regions of altered structural flexibility. Bottom: Volcano plot displaying $\log_2$ fold changes versus adjusted p-values for all structural fragment pairs. Significantly upregulated (red), downregulated (blue), and non-significant (gray) fragment pairs are distinguished. Significant fragments (FDR $< 10^{-4}$) reveal allosteric networks connecting position 203 to the substrate binding interface through specific structural segments.

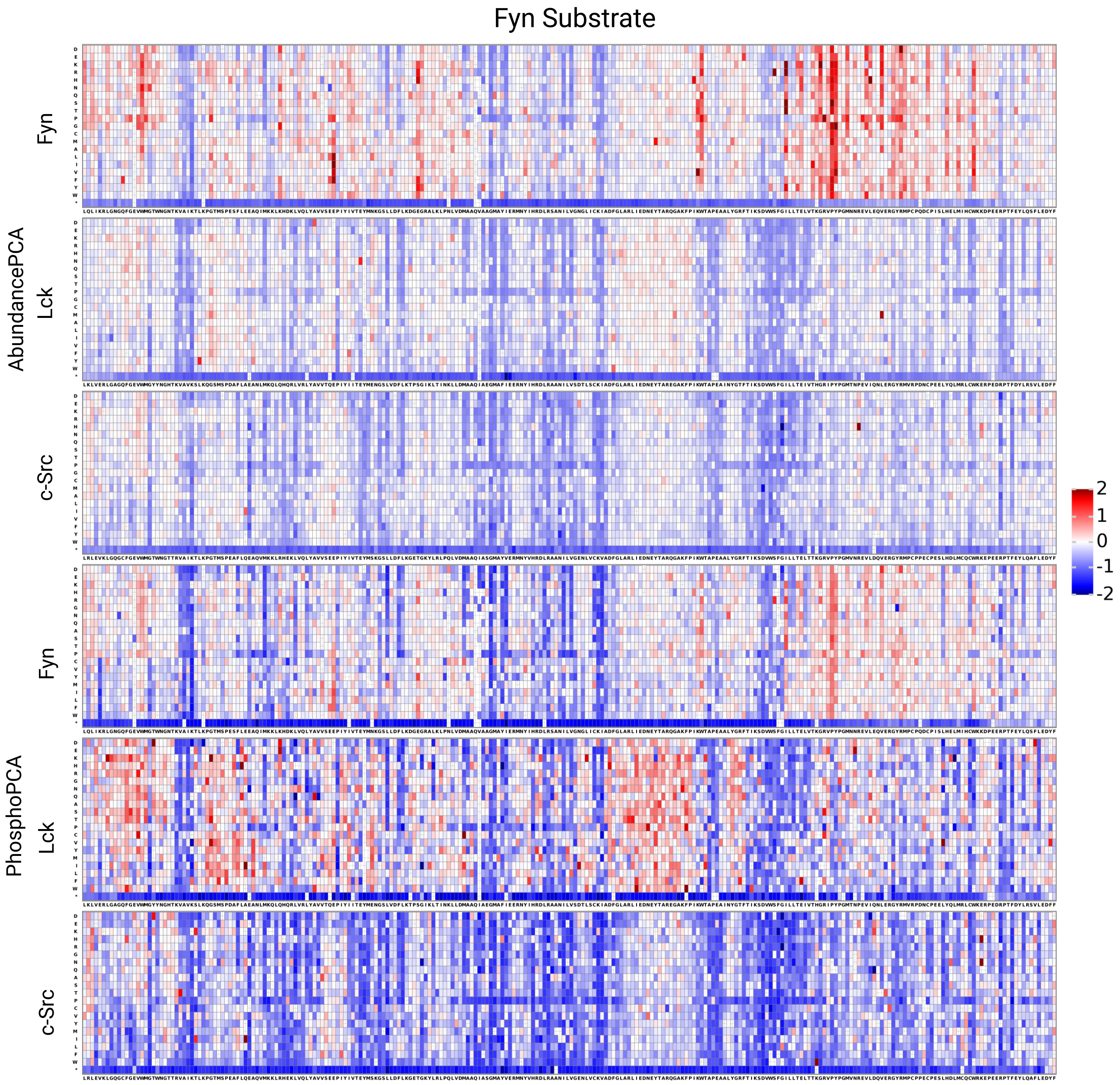


Figure S7: Complete mutational landscapes for all kinase-Fyn substrate pairs. Comprehensive heatmaps showing raw abundance-PCA (top three rows) and phospho-PCA (bottom three rows) scores for all 6 experimental conditions. Each panel displays scores for Fyn, Lck, or c-Src (rows) paired with Fyn-substrate. Colors indicate normalized fitness scores with red representing gain-of-function and blue representing loss-of-function effects. The 21 possible amino acid values (20 amino acids and stop codon) are shown on the y-axis, and the 254 positions along the kinase domain on the x-axis. These raw data serve as input for the hierarchical Bayesian decomposition analysis.

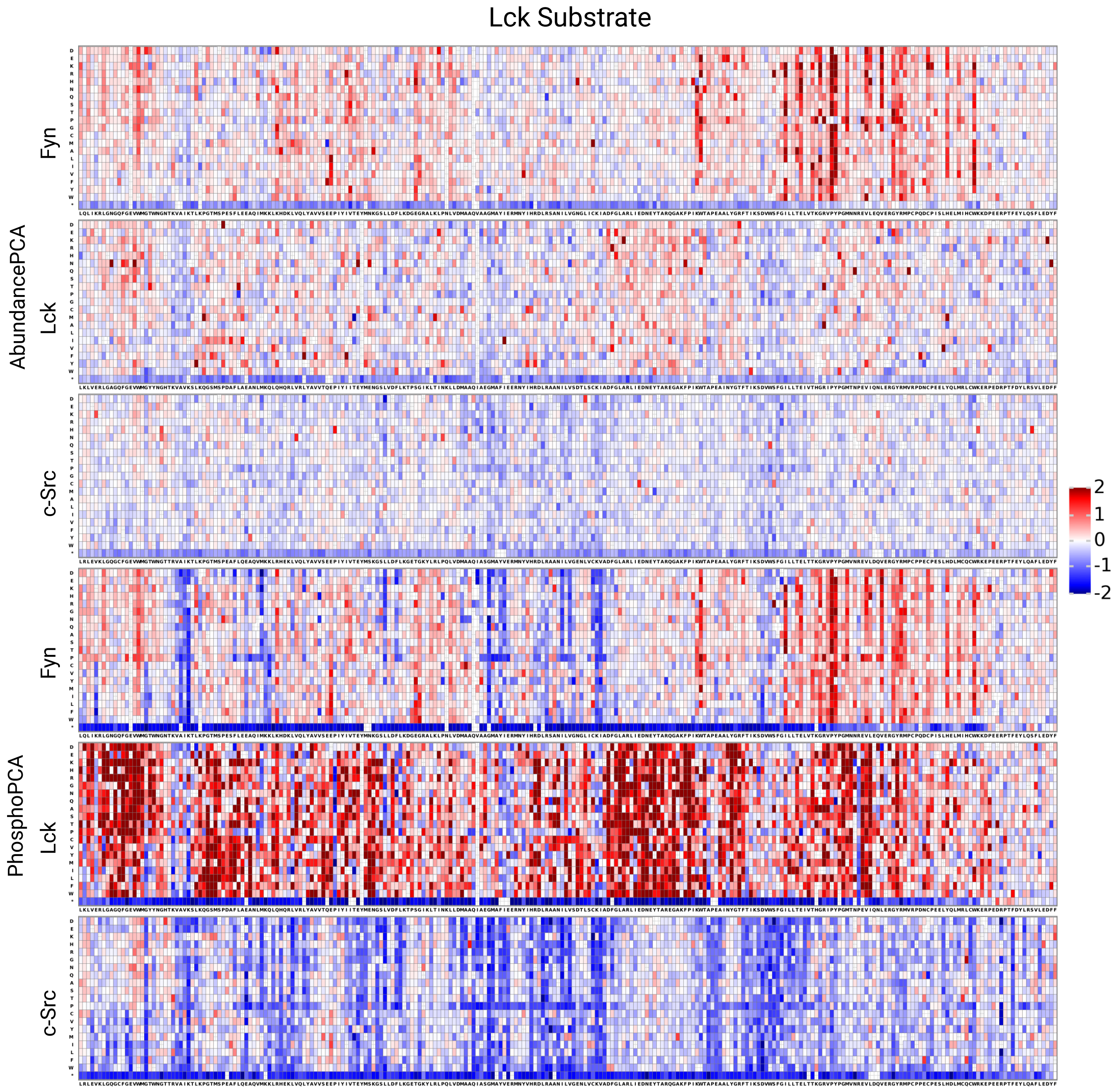


Figure S8: Complete mutational landscapes for all kinase-Lck substrate pairs. Comprehensive heatmaps showing raw abundance-PCA (top three rows) and phospho-PCA (bottom three rows) scores for all 6 experimental conditions. Each panel displays scores for Fyn, Lck, or c-Src (rows) paired with Lck-substrate. Colors indicate normalized fitness scores with red representing gain-of-function and blue representing loss-of-function effects. The 21 possible amino acid values (20 amino acids and stop codon) are shown on the y-axis, and the 254 positions along the kinase domain on the x-axis. These raw data serve as input for the hierarchical Bayesian decomposition analysis.

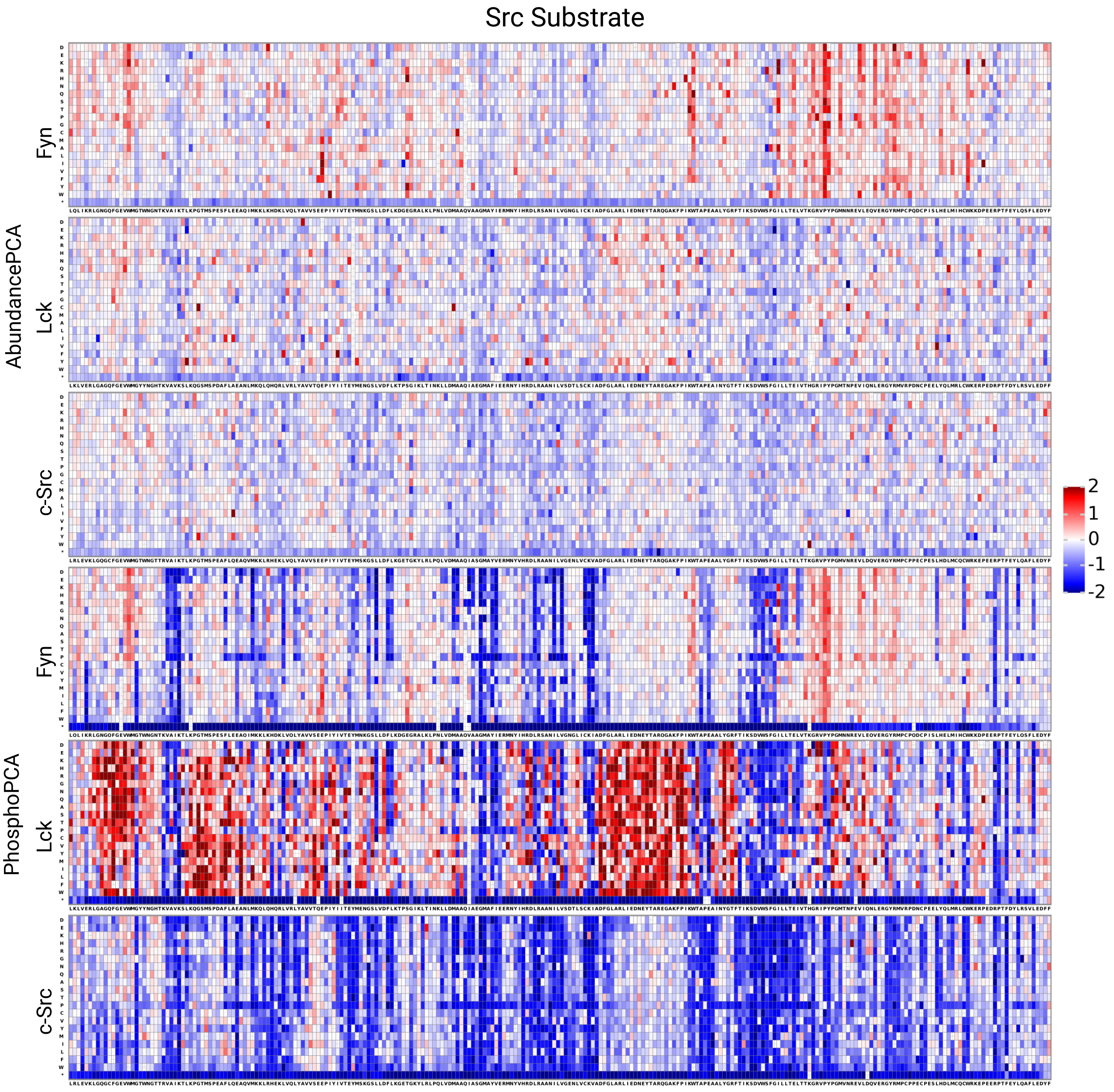


Figure S9: Complete mutational landscapes for all kinase-c-Src substrate pairs. Comprehensive heatmaps showing raw abundance-PCA (top three rows) and phospho-PCA (bottom three rows) scores for all 6 experimental conditions. Each panel displays scores for Fyn, Lck, or c-Src (rows) paired with c-Src-substrate. Colors indicate normalized fitness scores with red representing gain-of-function and blue representing loss-of-function effects. The 21 possible amino acid values (20 amino acids and stop codon) are shown on the y-axis, and the 254 positions along the kinase domain on the x-axis. These raw data serve as input for the hierarchical Bayesian decomposition analysis.

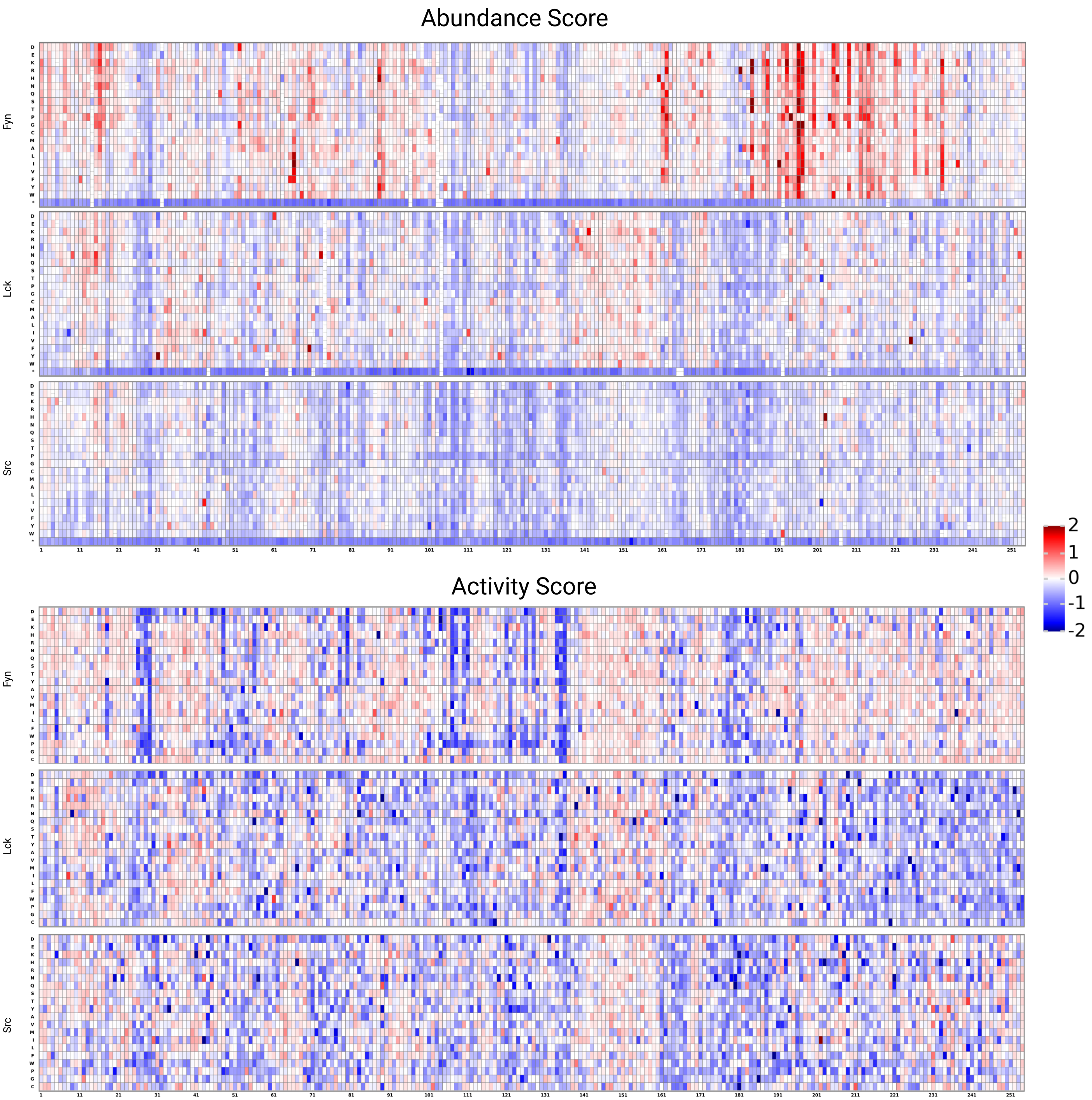


Figure S10: Pooled abundance and decomposed kinase activity scores across Src-family kinases. Top three panels: Mean of the abundance-PCA score shared across substrates for Fyn, Lck, and c-Src kinase domains, showing mutational effects on protein folding stability. Bottom three panels: Decomposed kinase activity scores capturing mutation-induced changes in catalytic function independent of abundance effects. These activity scores reflect conformational biasing and direct effects on catalysis, while removing confounding contributions from abundance. Positions with conserved loss-of-function or gain-of-function effects across kinases appear as columns of similar color.

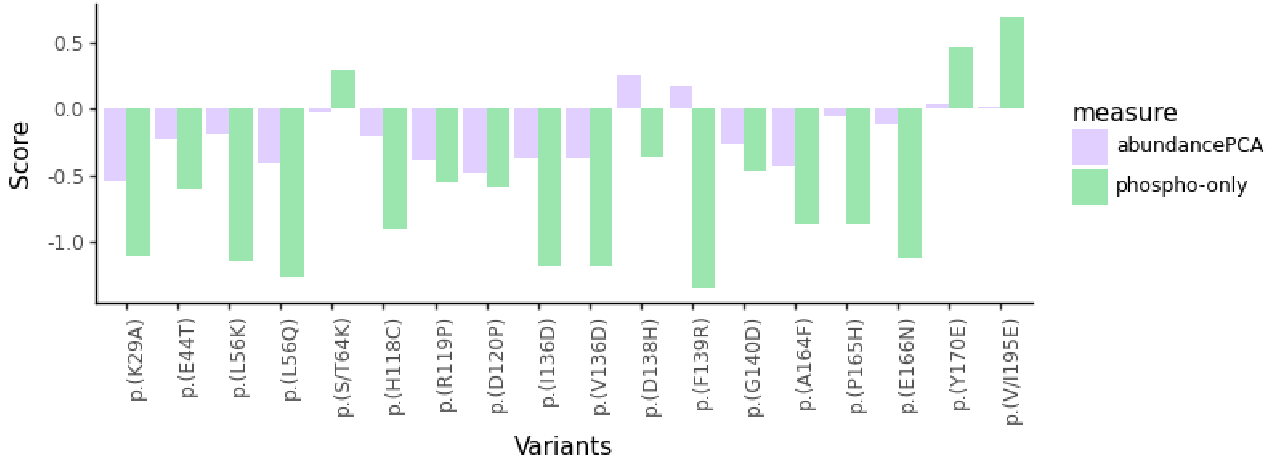


Figure S11: Abundance and decoupled activity scores identify catalytic motifs. Catalytic motifs (APE, HRD, DFG) are enriched for mutations with mild loss-of-function in abundance but severe defects in phosphorylation activity, yielding negative phospho-only scores across all three kinases. Residues p.(A164F), p.(P165H), p.(E166N) in the APE motif show strong phosphorylation dependence. The HRD motif variants p.(H118C), p.(R119P), p.(D120P) and DFG motif variants p.(D138H), p.(F139R), p.(G140D) exhibit mildly negative abundance scores but substantially more negative phosphorylation-only scores, with p.(F139R) showing the most conserved phospho-only loss-of-function across kinases. The previously uncharacterized p.(I/V136D), adjacent to the R-spine in the $Mg^{2+}$ positioning region, displays phosphorylation-only loss-of-function. Known catalytic sites at positions 29, 44, and 56 contain strongly negative activity scores, while positions 64, 170, and 195 harbor activity gain-of-function mutations.

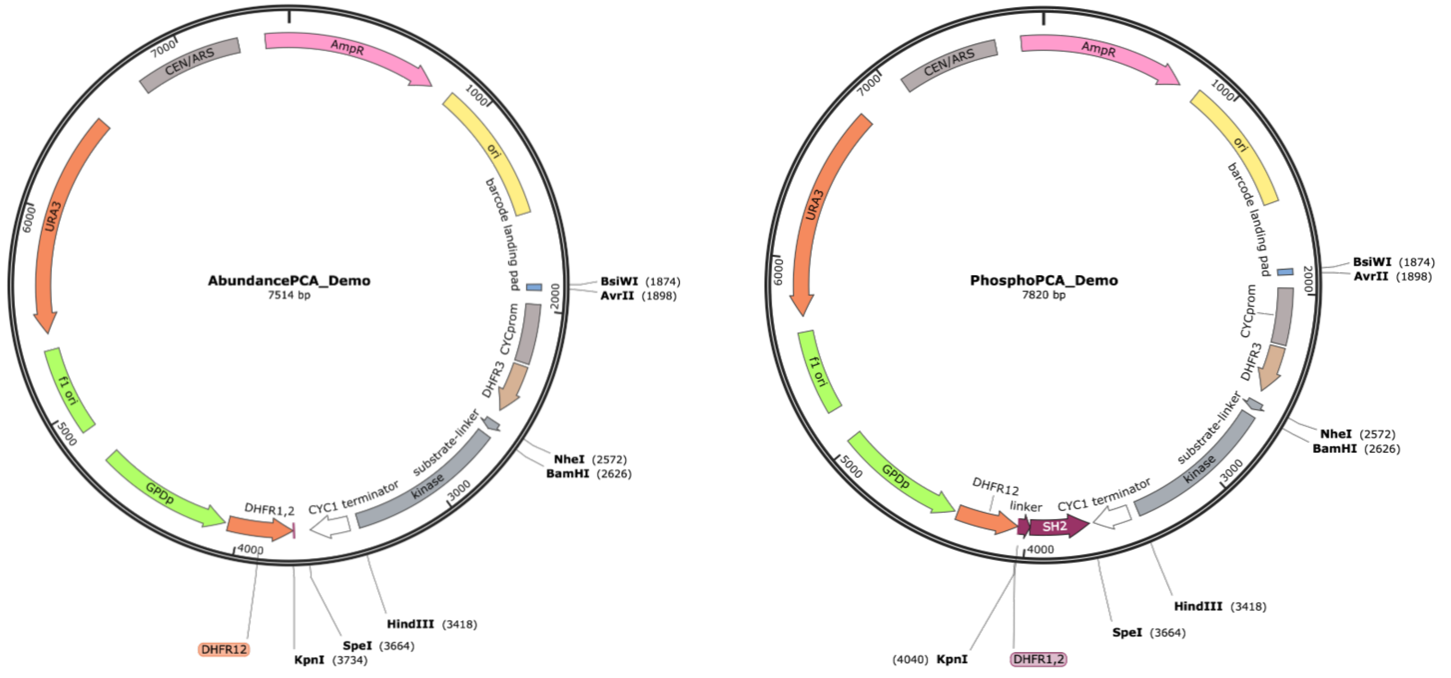


Figure S12: Plasmid maps for abundance-PCA and phospho-PCA plasmids.

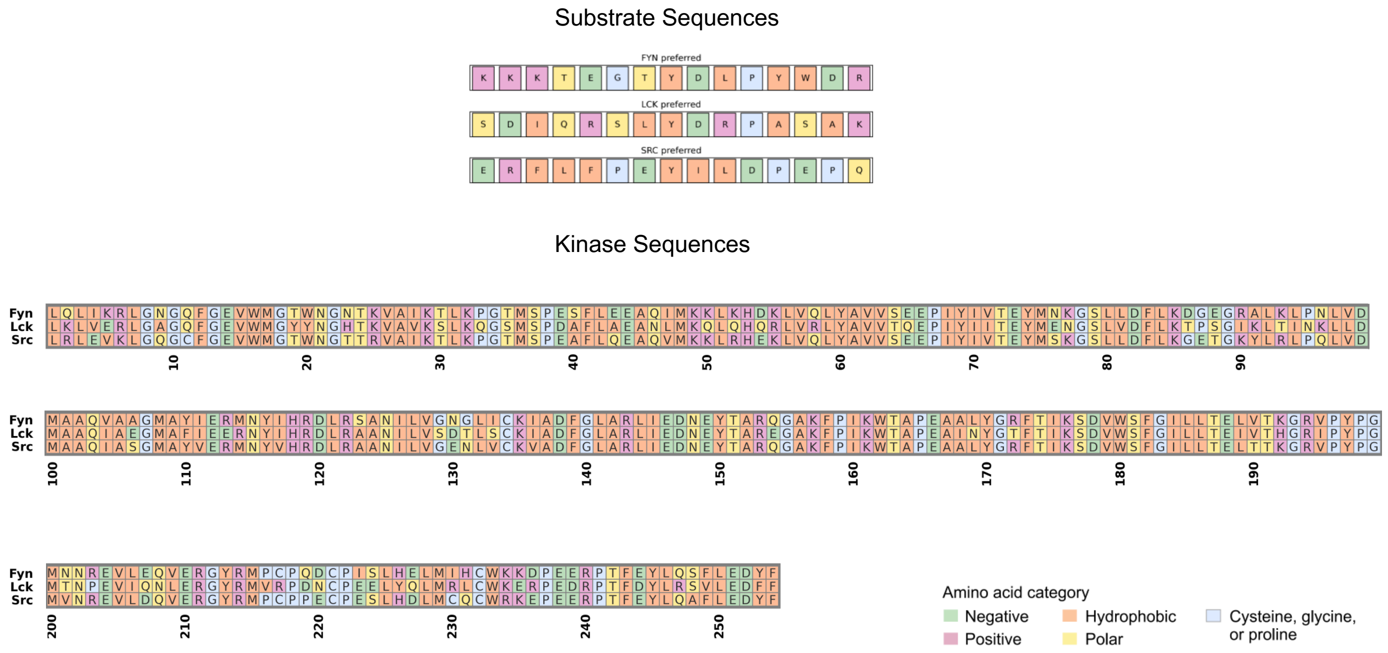


Figure S13: Sequences and residue-level chemical properties for Fyn, Lck, and c-Src and the respective substrates Fyn-sub, Lck-sub, and c-Src-sub.

| | lib_type | kinase | substrate | n_singlea_variants_barcodes | n_single_aa_variants | avg_barcode_coverage_single_aa | avg_variant_coverage_single_aa |
|---|---|---|---|---|---|---|---|
| 3 | PhosphoPCA | Fyn | lynsub | 53446 | 4902 | 12.126587 | 0.964961 |
| 4 | PhosphoPCA | Lck | lynsub | 49204 | 4898 | 10.045733 | 0.964473 |
| 5 | PhosphoPCA | Src | lynsub | 53495 | 4990 | 10.720441 | 0.982283 |
| 6 | AbundancePCA | Fyn | lynsub | 106737 | 4987 | 21.403046 | 0.981693 |
| 7 | AbundancePCA | Lck | lynsub | 82271 | 4988 | 16.493785 | 0.981890 |
| 8 | AbundancePCA | Src | lynsub | 99511 | 5046 | 19.228062 | 0.993307 |
| 12 | PhosphoPCA | Fyn | lcksub | 66269 | 4909 | 13.489491 | 0.966339 |
| 13 | PhosphoPCA | Lck | lcksub | 53772 | 4908 | 10.956090 | 0.966142 |
| 14 | PhosphoPCA | Src | lcksub | 62900 | 4992 | 12.600160 | 0.982677 |
| 15 | AbundancePCA | Fyn | lcksub | 61490 | 4910 | 12.523422 | 0.966539 |
| 16 | AbundancePCA | Lck | lcksub | 40235 | 4872 | 8.499043 | 0.959065 |
| 17 | AbundancePCA | Src | lcksub | 48966 | 4945 | 9.902123 | 0.973425 |
| 9 | PhosphoPCA | Fyn | srcsub | 124473 | 5001 | 24.888622 | 0.984449 |
| 10 | PhosphoPCA | Lck | srcsub | 83174 | 4987 | 16.679163 | 0.981693 |
| 11 | PhosphoPCA | Src | srcsub | 113412 | 5062 | 22.404583 | 0.996457 |
| 18 | AbundancePCA | Fyn | srcsub | 106458 | 4990 | 21.371706 | 0.980316 |
| 19 | AbundancePCA | Lck | srcsub | 67833 | 4962 | 13.670496 | 0.976772 |
| 20 | AbundancePCA | Src | srcsub | 86335 | 5039 | 17.133360 | 0.991929 |

Table S1: Sequencing coverage for abundance- and phospho-PCA libraries.

| Position | ss | Fyn interaction | Lck interaction | c-Src interaction | Fyn effect | Lck effect | c-Src effect |
|---|---|---|---|---|---|---|---|
| 28 | B3 | | | 26:CG2-28:CG1 | LOF | LOF | No Effect |
| 28 | B3 | 28:CG2-69:CG | 28:CG1-69:CG | | LoF | LoF | No Effect |
| 101 | HE | | 101:CB-186:CD2 | 101:CB-186:CD2 | No Effect | LoF | LoF |
| 101 | HE | | 101:O-182:CZ | 101:O-182:CZ | No Effect | LoF | LoF |
| 141 | ALN | 141:O-143:N | 141:CA-143:NH1 | | No Effect | No Effect | LoF |
| 147 | ALN | | 147:C-149:N | 147:C-149:N | LoF | No Effect | No Effect |
| 159 | ALC | | 122:NH1-159:CG | 122:NH1-159:CG | LoF | No Effect | No Effect |
| 159 | ALC | 120:OD1-159:CG | | | LoF | No Effect | No Effect |

Table S2: Divergent abundance-PCA mutations that disrupt van der Waals interaction networks. Residues are listed in multiple rows when they participate in more than one distinct interaction network.

| Position | Kinase | WT | Mutant | Fynsub | Lcksub | c-Srcsub |
|---|---|---|---|---|---|---|
| 9 | Lck | A | Q | 0.015 | −0.028 | 0.013 |
| 9 | Lck | A | S | 0.029 | −0.021 | −0.008 |
| 19 | Lck | Y | T | 0.004 | −0.026 | 0.022 |
| 19 | c-Src | T | A | −0.015 | 0.029 | −0.014 |
| 19 | c-Src | T | F | −0.003 | 0.026 | −0.022 |
| 33 | Lck | Q | P | 0.023 | −0.032 | 0.009 |
| 43 | Lck | A | S | 0.026 | −0.026 | 0 |
| 43 | Lck | A | T | 0.001 | −0.031 | 0.03 |
| 43 | c-Src | Q | E | 0.026 | −0.006 | −0.02 |
| 43 | c-Src | Q | V | −0.037 | 0.027 | 0.01 |
| 65 | Lck | Q | D | 0.034 | −0.029 | −0.005 |
| 90 | Lck | I | H | 0.023 | −0.026 | 0.003 |
| 113 | Lck | E | H | 0.021 | −0.036 | 0.015 |
| 113 | Lck | E | K | 0.014 | −0.025 | 0.011 |
| 114 | Lck | R | Y | 0.001 | −0.026 | 0.025 |
| 123 | Fyn | S | L | −0.025 | 0.015 | 0.009 |
| 172 | Lck | T | K | 0.013 | −0.025 | 0.012 |
| 172 | Lck | T | R | 0.026 | −0.017 | −0.008 |
| 203 | Lck | P | H | 0.019 | −0.027 | 0.008 |
| 203 | Lck | P | R | 0.024 | −0.026 | 0.002 |
| 226 | c-Src | H | W | 0.014 | 0.013 | −0.027 |
| 226 | c-Src | H | Y | −0.024 | 0.026 | −0.002 |
| 247 | Lck | R | Q | 0.009 | −0.026 | 0.017 |
| 248 | c-Src | A | N | 0.039 | −0.01 | −0.029 |
| 248 | c-Src | A | T | 0.032 | −0.016 | −0.016 |

Table S3: Specificity determining variants.

**Caption for Supplemental Data 1.** Short-read sequencing data. Raw short-read sequencing files available at https://doi.org/10.5281/zenodo.17541172.

**Caption for Supplemental Data 2.** Long-read sequencing data. Raw long-read sequencing files are available at https://doi.org/10.5281/zenodo.17541166.

**Caption for Supplemental Data 3.** Processed data and analysis pipelines. Processed datasets and computational analysis code are available at https://doi.org/10.5281/zenodo.17539481.

**Caption for Supplemental Movie 1.** Molecular dynamics simulations. Videos of molecular dynamics simulation trajectories are available at https://doi.org/10.5281/zenodo.17539481.
